# Solution-state NMR Assignment and Secondary Structural Propensities of the Full-Length and Minimalistic-Truncated Prefibrillar Monomeric Form of Biofilm-Forming Functional-Amyloid FapC from *Pseudomonas aeruginosa*

**DOI:** 10.1101/2023.01.22.525107

**Authors:** Chang-Hyeock Byeon, Pang C. Wang, In-Ja L. Byeon, Ümit Akbey

**Affiliations:** Department of Structural Biology, School of Medicine, University of Pittsburgh, Biomedical Science Tower 3, 3501 Fifth Avenue, Pittsburgh, 15261, United Stated

**Keywords:** Solution-state NMR Spectroscopy, functional bacterial amyloid (FuBA), bacterial biofilm, *Pseudomonas aeruginosa* FapC, structure determination, prefibrillar assembly

## Abstract

Functional bacterial amyloids provide structural scaffolding to bacterial biofilms. In contrast to the pathological amyloids, they have a role *in vivo* and are tightly regulated. Their presence is essential to the integrity of the bacterial communities surviving in biofilms and may cause serious health complications. Targeting amyloids in biofilms could be a novel approach to prevent chronic infections. However, structural information is very scarce on them in both soluble monomeric and insoluble fibrillar forms, hindering our molecular understanding and strategies to fight biofilm related diseases. Here, we present solution-state NMR assignment of 250 amino acid long biofilm-forming functional-amyloid FapC from *Pseudomonas aeruginosa*. We studied the full-length and shorter minimalistic-truncated FapC constructs without signal-sequence that is required for secretion. 91% and 100% backbone NH resonance assignment for FL and short constructs, respectively, indicates that soluble monomeric FapC is predominantly disordered, with sizeable secondary structural propensities mostly as PP2 helices, but also as α-helices and β-sheets highlighting hotspots for fibrillation initiation interface. Shorter construct showing almost identical NMR chemical shifts highlights the promise of utilizing it for more demanding solid-state NMR studies that requires methods to alleviate signal redundancy due to almost identical repeat units. This study provides key NMR resonance assignment for future structural studies of soluble, pre-fibrillar and fibrillar forms of FapC.

## Biological Context

FapC is the major biofilm forming amyloid fibril from *Pseudomonas aeruginosa*.^1, 2^ FapC does not directly cause disease in contrast to the pathological amyloid fibrils and has a unique function in the biofilm. It thus belongs to the emerging group of functional bacterial amyloids (FuBA), such as CsgA, TasA and many others.^2-8^ Along with its minor components it constitutes the structural scaffold of bacterial colonial biofilms. Biofilms act as a shield for bacteria and protects them from external effects such as antibiotics, and cause most of the persistence chronic infections. When functional amyloids are mutated, the biofilm loses its rigidity, and the bacteria becomes accessible.^4^ As a result, antibiofilm agents are a promising source and research avenues to fight antimicrobial resistance. Structural insights provided here by NMR will be essential in aiding the molecular understanding of functional amyloid formation, its interactions with other protein and biofilm components, as well as to design better molecules targeting amyloids and biofilms.

Wild-type FapC is a 250 amino acid (aa) protein with a signal peptide (24aa) that targets the protein to outside the bacteria. The functional 226 amino acid secreted protein is composed of an N-terminus (37 aa) and a C-terminus (21 aa), three almost identical repeats (3*30 aa) separated by two longer flexible loops (L1: 35 and L2: 43 aa), Figure 1A. This structural construction differs from CsgA that consists of five consecutive shorter repeats of ∼20 aa and from TasA that consists of a non-repeat based longer protein sequence of 233 aa. FapC and CsgA are unfolded intrinsically disordered proteins (IDP).^5^ TasA, however, has a globular folded protein fold.^5, 9^ FapC has been predominantly characterized by biophysical techniques and low-resolution methods as a pure protein or with its interaction partners. A high-resolution NMR based study with sequential signal assignments was missing. We reported a solid-state NMR (ssNMR) based initial study on FapC in its solid fibrillar form.^10^ In this work, we demonstrate a divide and concur approach for NMR based structure determination protocols, since the almost identical three repeats impose a dramatic challenge particularly for ssNMR based signal assignment. As a result, a full-length (FL; NR1L1R2L2R3C, N: N terminus, R: repeat, L: loop, and C: C terminus) and a shorter truncated L2R3C FapC constructs were prepared. From these shorter constructs, the minimalistic truncated L2R3C were prepared that contains singular key structural elements as one repeat unit, one loop and the C-terminus with a total of 94 amino acids. Here, finally we present the solution-state NMR signal assignment of L2R3C construct and compare its spectra with the FL FapC. A complete NMR signal assignment is elaborated by a secondary structure propensity calculation. It came to our attention at the time of our submission that another preprint shows the solution-state NMR signal assignment of FL FapC with different sample conditions with additives such as a chaperone.^11^ No BMRB data was available at the time of our submission, and we present results solely based on our own manual NMR signal assignment work performed last year and the data is deposited to the BMRB database.

**Figure 1.**
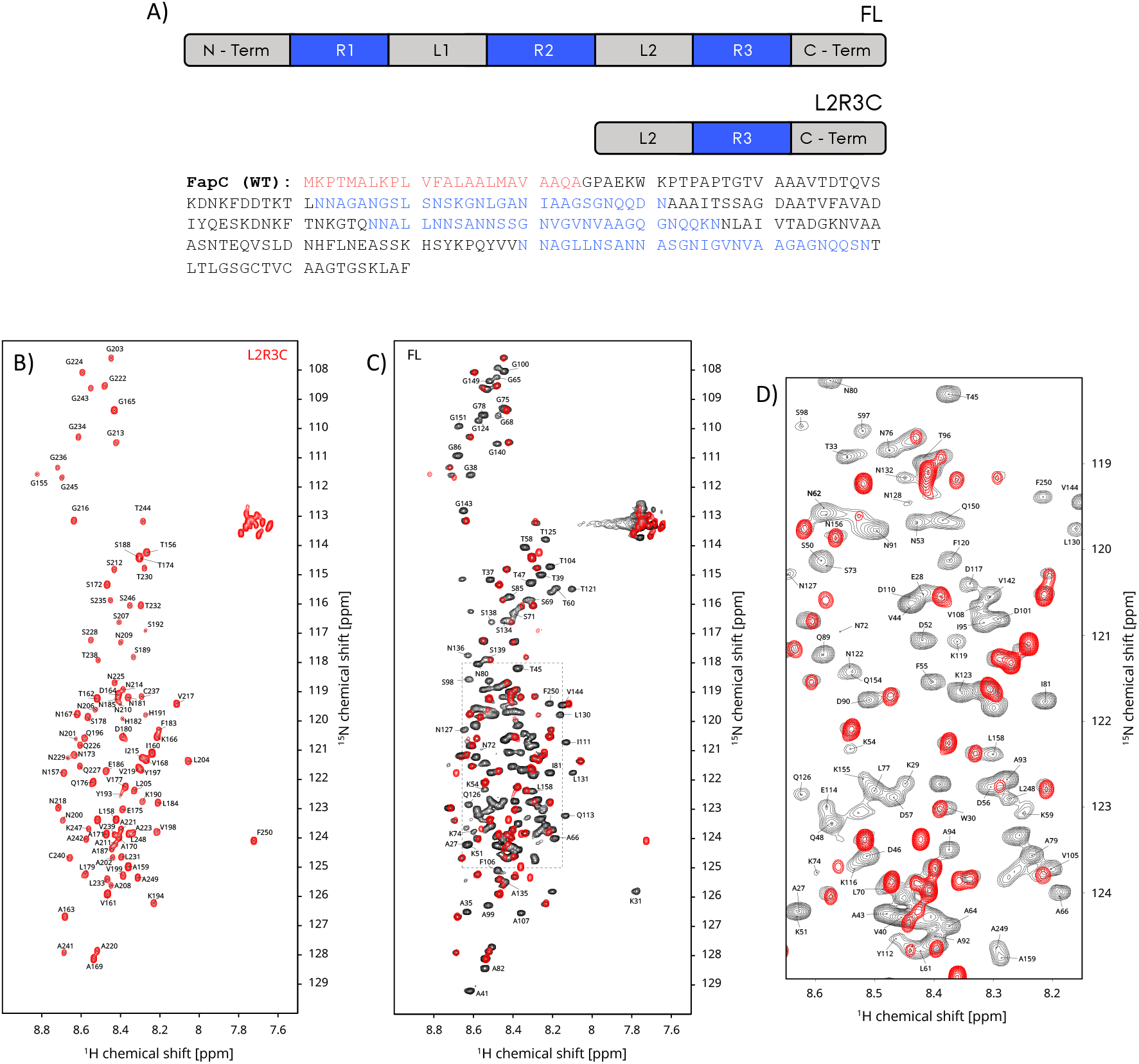
**A)** Schematic domain representation of the FL (residues 25-250, 226 aa, no signal peptide) and L2R3C (residues 157-250, 94 aa) FapC constructs. The amino acid sequence is given as well with color coding matching the representation. ^1^H-^15^N HSQC NMR spectra of FL (black) and L2R3C (red) FapC samples. **B)** The signal assignment for L2R3C FapC sample. **C)** The overlay of the L2R3C and FL NMR spectra. Note that the spectra are well overlapped, indication of structural similarity. Residues outside the highlighted area that are well separated are shown, only for the FL specific residues that do not exist in L2R3C. **D)** The zoomed area for the crowded highlighted region in C. Only the resonance assignments from the FL (black) spectrum are shown. For the L2R3C spectral assignments (in red) in A.

### Recombinant protein production

The FL construct (residues 25-250, 226 aa, no signal peptide) was cloned into a pET28a vector using NcoI and XhoI, and it expressed the FL protein containing nine additional amino acids due to the addition of one methionine on the N-terminus and histidine purification tag (LEHHHHHH) at the C-terminus. Recombinant FapC L2R3C (residues 157-250, 94aa) plasmid was cloned into pET32 vector using the Takara Bio In-fusion cloning mechanism. A modified protocol was used to produce the L2R3C construct to increase the yield of the expressed protein. L2R3C was produced from the appropriate FL FapC construct with additional Thioredoxin (Trx), 10 histidine (His10) tag, and thrombin cleavage site at the N-terminal. The final cleaved L2R3C protein contained four additional amino acids at the N-terminus for cloning and cleavage artifacts (GSGT).

Bacterial plasmids were transformed into the BL21(DE3) E. coli bacteria. All nonisotope-labeled samples were prepared in LB media. Isotope labeling for NMR studies was carried out by growing in minimal media with ^13^C-glucose and ^15^N-ammonium chloride as carbon and nitrogen sources, respectively. The transformed BL21(DE3) cells were plated onto LB agar with the appropriate antibiotic and grown overnight in 37 °C incubator. The colonies were resuspended in the media listed above with appropriate antibiotic and inoculated into large volume media. These cultures were grown in shaker incubator at 37 °C until the OD600 reached a value between 0.8-1.0 OD. For induction, IPTG was added to each culture for a final concentration of 1 mM and grown for another 4 hours at 37 °C. Cells were harvested by centrifugation (6000 RCF, for 20 min at 4 °C). The cells were then lysed using a sonicator in denaturing buffer (50 mM Tris and 8 M guanidinium chloride (GdmCl), pH 8.0. Purification was carried out on a His-Tag affinity column by step-eluting the protein in 30-500 mM imidazole (in addition to 50 mM Tris pH 8 and 8 M GdmCl). FapC containing fractions were pooled together. FapC FL was spun down (20800 RCF, 10 min at RT) and the FL containing supernatant was stored at -20 °C.

FapC L2R3C protein was produced with additional cleave and purification steps. The tagged L2R3C protein was desalted into 50 mM sodium phosphate, pH 7.4 buffer in PD-10 column (GE Healthcare) using Sephadex G-25 medium according to the manufacturer’s protocol. Thrombin protease (Thermo Scientific) was added at 4U per mg of tagged protein and digested at 4 °C for 16 hours. The complete cleavage was verified by SDS-PAGE and LC-MS (Thermo Fisher), then solid GdmCl was added to the solution to reach a final concentration of 8 M. The Trx-His10-tag was separated from the cleaved L2R3C protein with His-

Tag affinity column (in the presence of 50 mM Sodium Phosphate and 8 M GdmCl, pH 7.4) by trapping the tag at the column, while washing and collecting the L2R3C in the flow through. The FapC L2R3C was concentrated and stored at room temperature. All proteins were desalted into 50 mM sodium phosphate, pH 7.4 buffer on a PD-10 column (GE Healthcare) using Sephadex G-25 medium prior to making the NMR samples. ∼60 and ∼20 mg of uniformly ^13^C,^15^N isotope labeled pure protein were obtained per liter of culture for FL and L2R3C, respectively.

### NMR Spectroscopy

NMR spectra of uniformly ^13^C,^15^N isotope labeled FapC FL (226 aa, without the signal peptide) and L2R3C (94 aa) were recorded at Bruker Avance III 600, Avance 700 and Avance II 900 MHz spectrometers equipped with a 5 mm triple resonance TCI cryoprobes. We used 5 mm NMR sample tubes with a total volume of 500 μl and a protein concentration of ∼0.3 mM in 50 mM sodium phosphate, 30mM DTT, 1 mM D6-DSS, 10% D_2_O and pH 7.4 buffer. The NMR samples were prepared freshly prior performing the 3D NMR experiment blocks. All NMR experiments were recorded at 274 K. The details of the NMR experiments are shown in **Table 1**. 2D ^1^H-^15^N HSQC and 3D HNCACB, HN(CO)CACB spectra were used for C_α_ and C_β_ assignments. 3D CC(CO)NH TOCSY, H(CCCO)NH TOCSY and HBHA(CO)NH spectra were used for sidechain assignments. The mixing time was 20 ms for the TOCSY experiments. We also recorded time-shared 3D ^13^C-editted/^15^N-editted NOESY HSQC spectra to further identify intra and inter residue connectivities. NOESY mixing time was 200 ms. The spectra were processed by Topspin 3 (Bruker Biospin), except for the time-shared 3D NOESY experiment which was processed by NMRPipe.^12^ The NMR resonance assignment was done manually by using CCPNmr Analysis 3.^13^ The ^1^H chemical shifts were referenced to 0 ppm by using DSS as an external standard added to the NMR samples. The ^13^C and ^15^N chemical shifts were indirectly referenced by using the ^1^H frequency as explained previously.^14^

**Table 1.** The experimental conditions of the 2D and 3D NMR experiments performed in this study. The spectra were recorded with conventional incrementation of the indirect dimensions. The spectra were recorded at 274K.

| NMR Experiment | ns | RD | TD direct | Increment | SW | TD | Increment | SW | TD | Increment | SW | TD | Increment | SW | Instrument | Expt. Time |
| --- | --- | --- | --- | --- | --- | --- | --- | --- | --- | --- | --- | --- | --- | --- | --- | --- |
| | | s | 1H | μs | ppm | 15N | μs | ppm | 13C | μs | ppm | 1H | μs | ppm | MHz | hour |
| 2D HN HSQC | 8 | 1.0 | 2k | 39.6 | 14 | 256 | 456.7 | 24 | - | - | - | - | - | - | 600/900 | 1 |
| 3D HNCACB | 64 | 1.0 | 2k | 39.6 | 14 | 50 | 456.7 | 24 | 80 | 73.6 | 60 | - | - | - | 900 | 72 |
| 3D HN(CO)CACB | 64 | 1.0 | 2k | 39.6 | 14 | 46 | 498.2 | 22 | 80 | 73.6 | 60 | - | - | - | 900 | 72 |
| 3D CC(CO)NH TOCSY | 128 | 0.9 | 2k | 44.4 | 16 | 36 | 587.4 | 24 | 36 | 81.2 | 70 | - | - | - | 700 | 42 |
| 3D H(CCCO)NH TOCSY | 128 | 0.9 | 2k | 44.4 | 16 | 36 | 587.4 | 24 | - | - | - | 36 | 238.1 | 6 | 700 | 42 |
| 3D HBHA(CO)NH | 128 | 0.9 | 2k | 44.4 | 16 | 36 | 587.4 | 24 | - | - | - | 36 | 238.1 | 6 | 700 | 42 |
| 3D 13C/15N edited NOESY | 128 | 1.0 | 2k | 30.8 | 18 | 32 | 456.7 | 24 | 32 | 184.1 | 24 | 108 | 92.6 | 12 | 900 | 123 |

### NMR Assignment, Statistics and Data Deposition

The 2D ^1^H-^15^N HSQC spectra of the L2R3C and FL samples are shown in Figure 1. The NMR spectrum of the L2R3C sample (red spectrum in B) is of very high quality due to the favorable properties of the disordered protein, despite small chemical shift dispersion. The backbone NH resonances are completely assigned excluding one proline (P195), shown in Figure 1B. Here is the summary of overall FapC resonance assignment statistics for L2R3C (94 aa): 100% backbone HN nuclei (93/93 excluding one proline), 100% of the C_α_ (94/94), 100% of the C_β_ (84/84), 96% of the H_α_ protons (90/94), 95% of the H_β_ protons (80/84), and FL FapC (226 aa): 91% backbone HN nuclei (201/221 excluding five proline), 97% of the C_α_ (220/226), 98% of the C_β_ (198/202), 88% of the H_α_ protons (196/224), 91% of the H_β_ protons (184/202). By using the H(CCCO)NH TOCSY spectrum recorded on L2R3C, we assigned 61% sidechain non-H_α_/H_β_ protons (89 out of 146 resonances). All the chemical shifts have been deposited in BMRB. A representative sequential NMR assignment stretch is shown in Figure 3 for residues 187-194 (A187-S188-S189-K190-H191-S192-Y193-K194) based on the 3D HNCACB spectra. Resonances from the i (residue itself) and i-1 (preceding) residues are labeled accordingly. The signal overlap in the carbon dimension of the 3D spectra is extremely challenging for some residues in the FL spectra, and the resonances could hardly be assigned unambiguously without the L2R3C spectra.

By using these C_α_ C_β_ NMR chemical shifts (Δδ(C_β_)-Δδ(C_β_)), we determined the deviation from random coil chemical shift values to identify stable α-helices or β-sheets.^18^ These values fall below the 1.4 ppm threshold (between ±1.1 ppm, data not shown), an indication of the IDP nature of the protein with random coil conformation with small structural propensities. Two different algorithms were used for secondary structure propensities (SSP), the neighbor-corrected secondary structure propensity (ncSSP) based on the work by Tamiola *et al*. and the δ2D method based SSP based on the work by Camilloni *et al*.^15,16,17^ Both methods are optimized for IDPs and resulted in similar overall structural propensities that indicate the probability of the protein to be in α-helix, β-sheet, PP2 helix or random coil conformation for similar regions of FapC (Figure 2). For the L2R3C construct, the residues between 181-190 indicates an opposite trend compared to the rest of the protein, indicating an α-helical conformational at ∼15% propensity compared to the remaining β-sheet propensities of ∼5-25% with a few irregularities, Figure 2A-B. These patterns are similarly observed for the FL FapC, Figure 2C-D. The residues between 25-157 (the anti-L2R3C region in FL FapC) have alternating SSP, surprisingly with a large β-sheet propensity at the N-terminus. Overall, the NMR derived SSPs fit to the amyloid formation propensity predictions we reported previously.^10^

**Figure 2.**
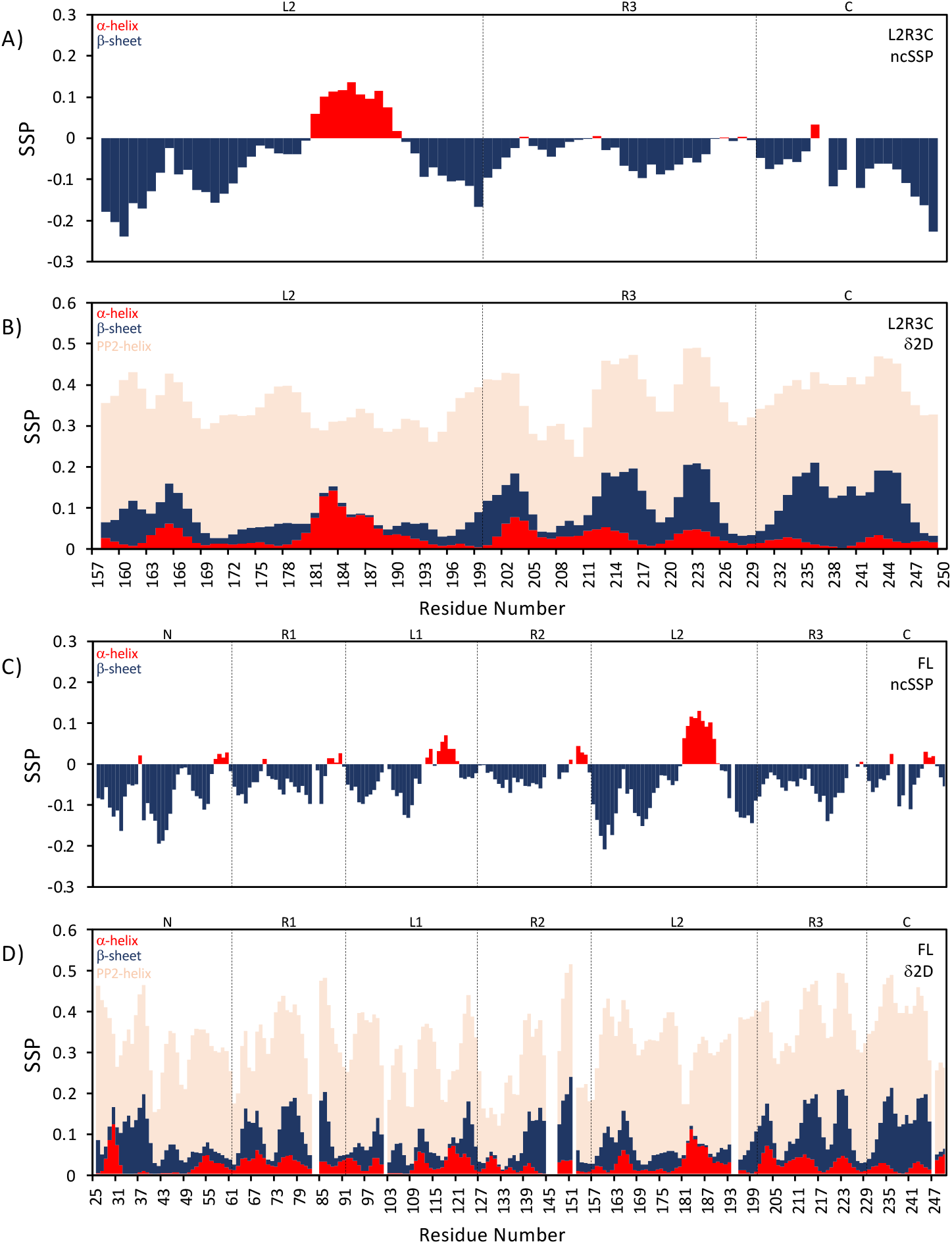
Residue-specific NMR chemical shift based secondary structure propensities (SSP) for α-helix (red), β-sheet (blue) and polyproline-2 (PP2) helix (pink) based on two different algorithms. A and B show results for L2R3C, and C and D for FL FapC. **A)** and **C)** The neighbor-corrected secondary structure propensities (ncSSP) based on the work by Tamiola *et al*.^15, 16^ **B)** and **D)** The δ2D method based secondary structure propensities based on the work by Camilloni *et al*.^17^ The rest of the propensities in B and D are random coil to complete the sum to 1.0, which is not shown explicitly. The propensities are shown in terms of percentage, whereby 1.0 equal to 100% propensity. The proposed secondary structure elements (based on amyloidogenic regions and sequence of FapC) are shown at each graph to guide the eye.

**Figure 3.**
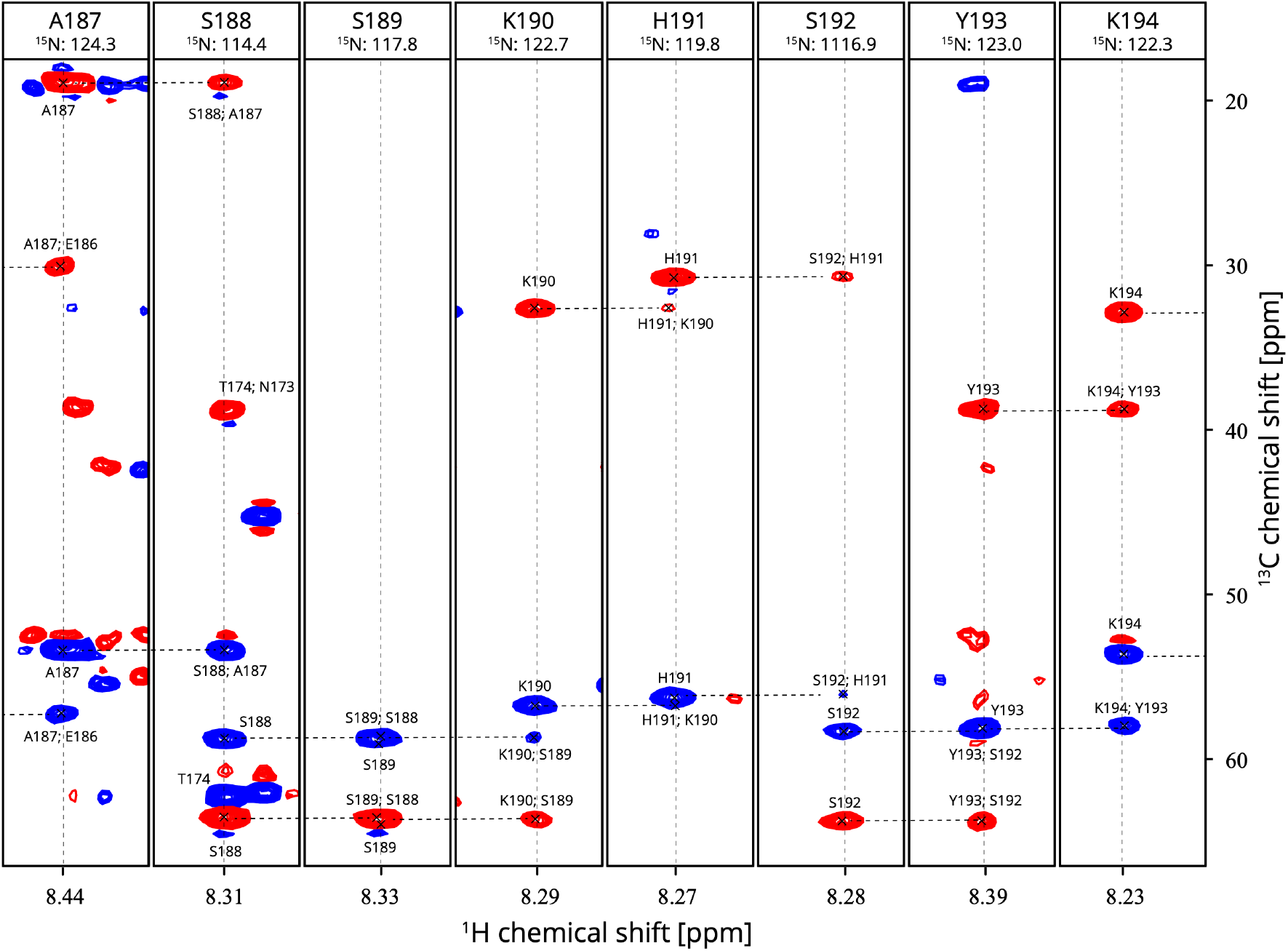
Representation of the NMR assignment for the L2R3C FapC sample for residues 187-194 (A187-S188-S189-K190-H191-S192-Y193-K194). The 3D HNCACB NMR spectrum is shown for this representation. The C_β_ and C_β_ resonances are in blue (positive signals) and red (negative signals) colors, respectively. Dashed lines are shown to guide the eye for the sequential connectivities between neighboring residues. The center of each strip is marked with a dashed line, and the corresponding proton and nitrogen chemical shifts are given in ppm. Signals from the i (residue itself) and i-1 (preceding) residues are labeled accordingly.

## Conclusions

In summary, we provide backbone proton and nitrogen resonance assignments 91% for the FL (NR1L1R2L2R3C) and 100% for the L2R3C minimalist FapC in their monomeric soluble prefibrillar form, excluding the backbone prolines. We compared the L2R3C and FL NMR spectra, showing no chemical shift changes between two protein forms, except the few beginning and end residues of the two constructs. The NMR chemical shift based SSP analysis reveals that the L2R3C FapC adopts most of the time a random coil or PP2-helix. Nevertheless, a distinct α-helical conformational propensity exists for L2R3C for residues between 181-190 along with the remaining β-sheet propensities at ∼20%. Remarkably, the α-helical and β-sheet SSP of the L2R3C region is preserved in the FL FapC construct. This study presents a first important step towards more detailed structural studies of FapC and presents the feasibility of the reduced complexity approach towards native-like structure determination protocols.

## Data Availability

NMR assignments been deposited in BMRB for L2R3C (#51792) and FL (#51793) FapC, release date is asap. The list of the NMR chemical shifts for L2R3C and FL FapC is included in the Supplementary Information. Data will be available upon request.

## Author Contribution

Conceptualization: UA; Methodology: CHB, IJLB, UA; Formal analysis and investigation: CHB, PCW, IJLB, UA; Writing - original draft preparation: CHB, UA; Funding acquisition: UA; Resources: UA; Supervision: UA.

## Acknowledgement

We acknowledge Jasper Jeffrey, Kasper H. Hansen, Maria Andreasen, Andy Hinck and Angela Gronenborn for helpful discussions. UA acknowledges financial support from University of Pittsburgh startup funding and the high-field NMR infrastructure at the Structural Biology Department, School of Medicine, University of Pittsburgh.

## Supplementary Information

**Table 1.**
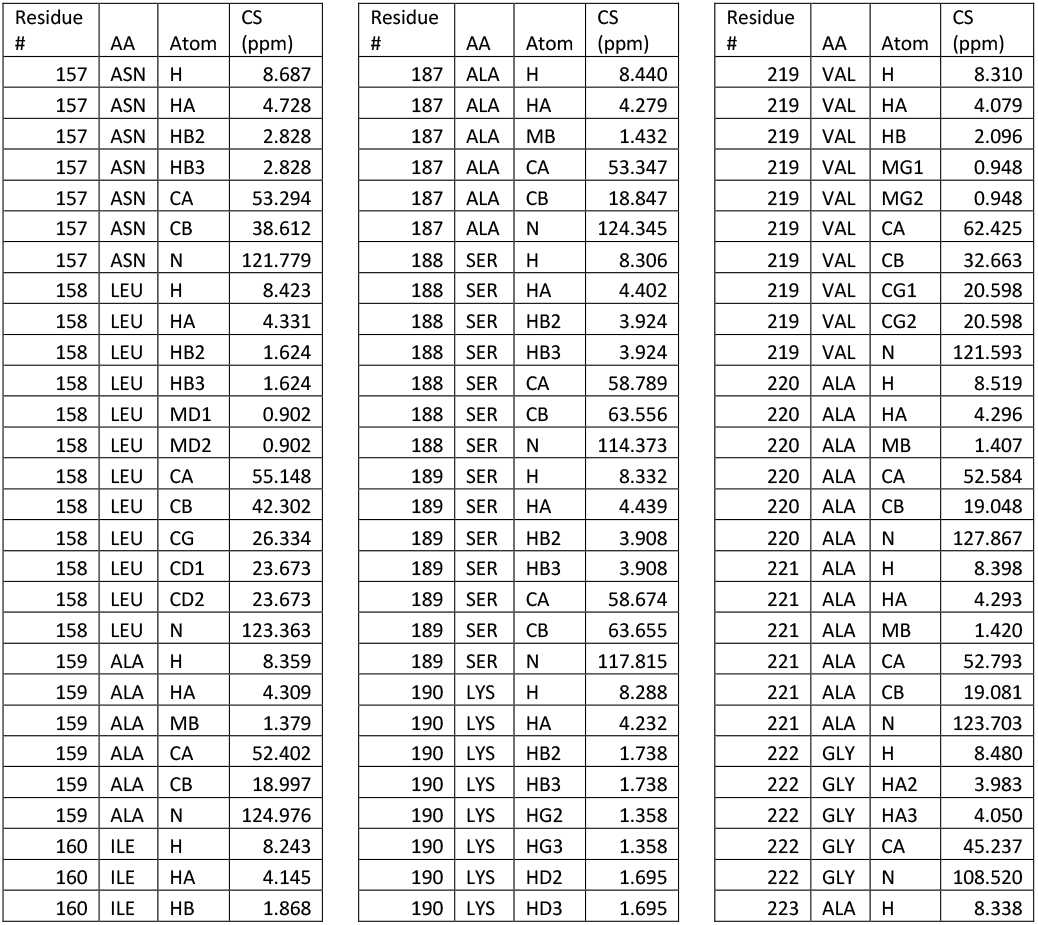
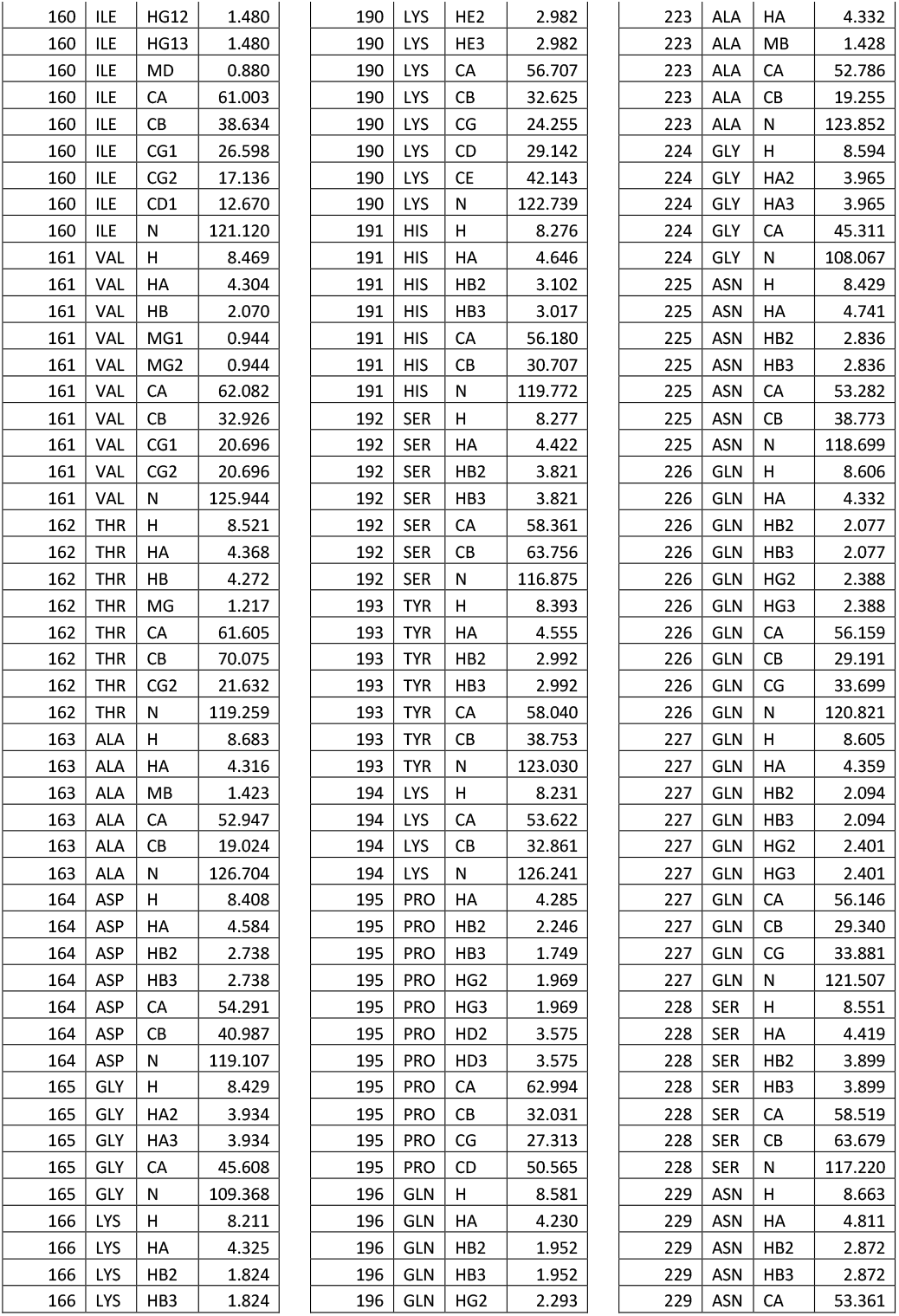
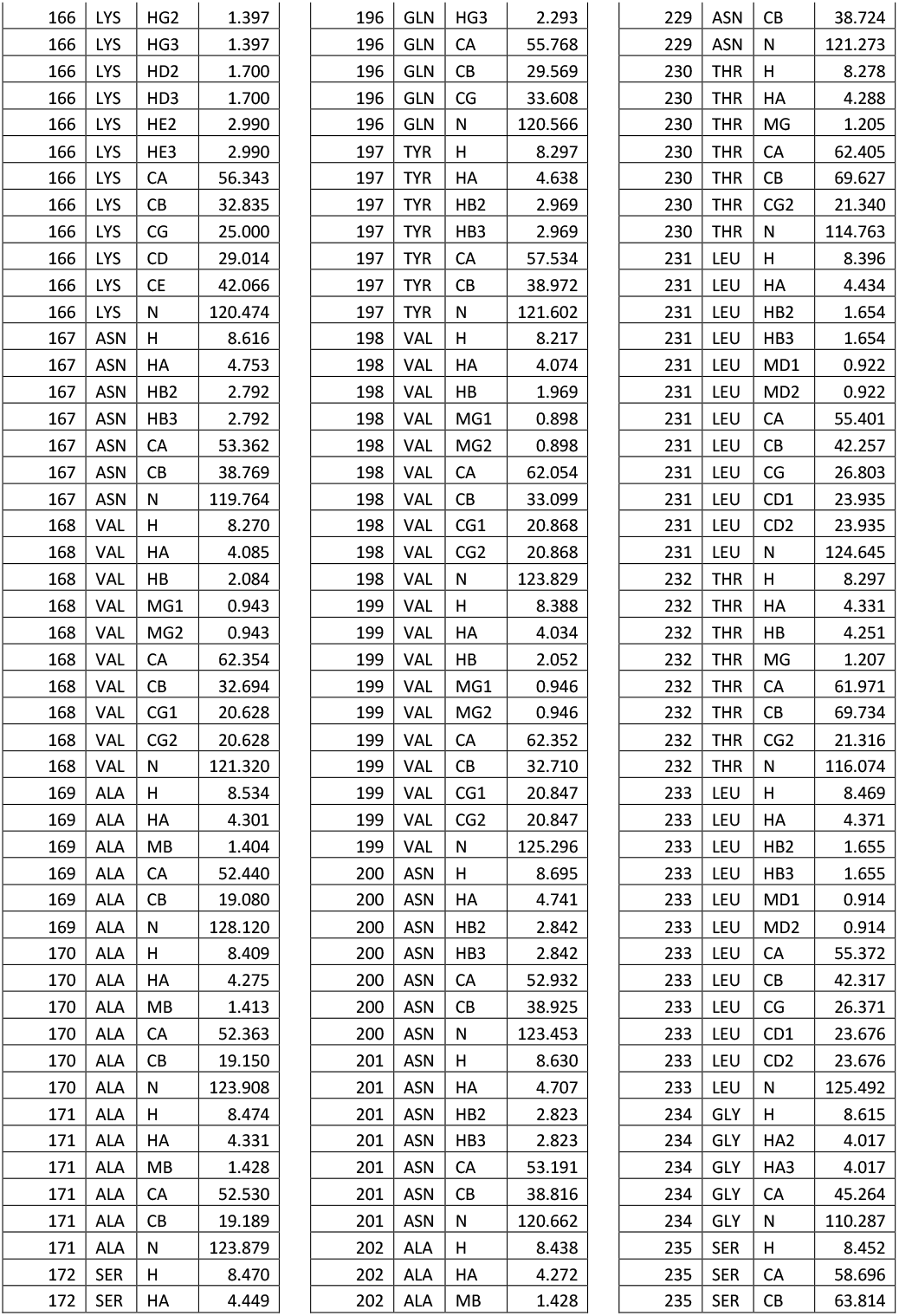
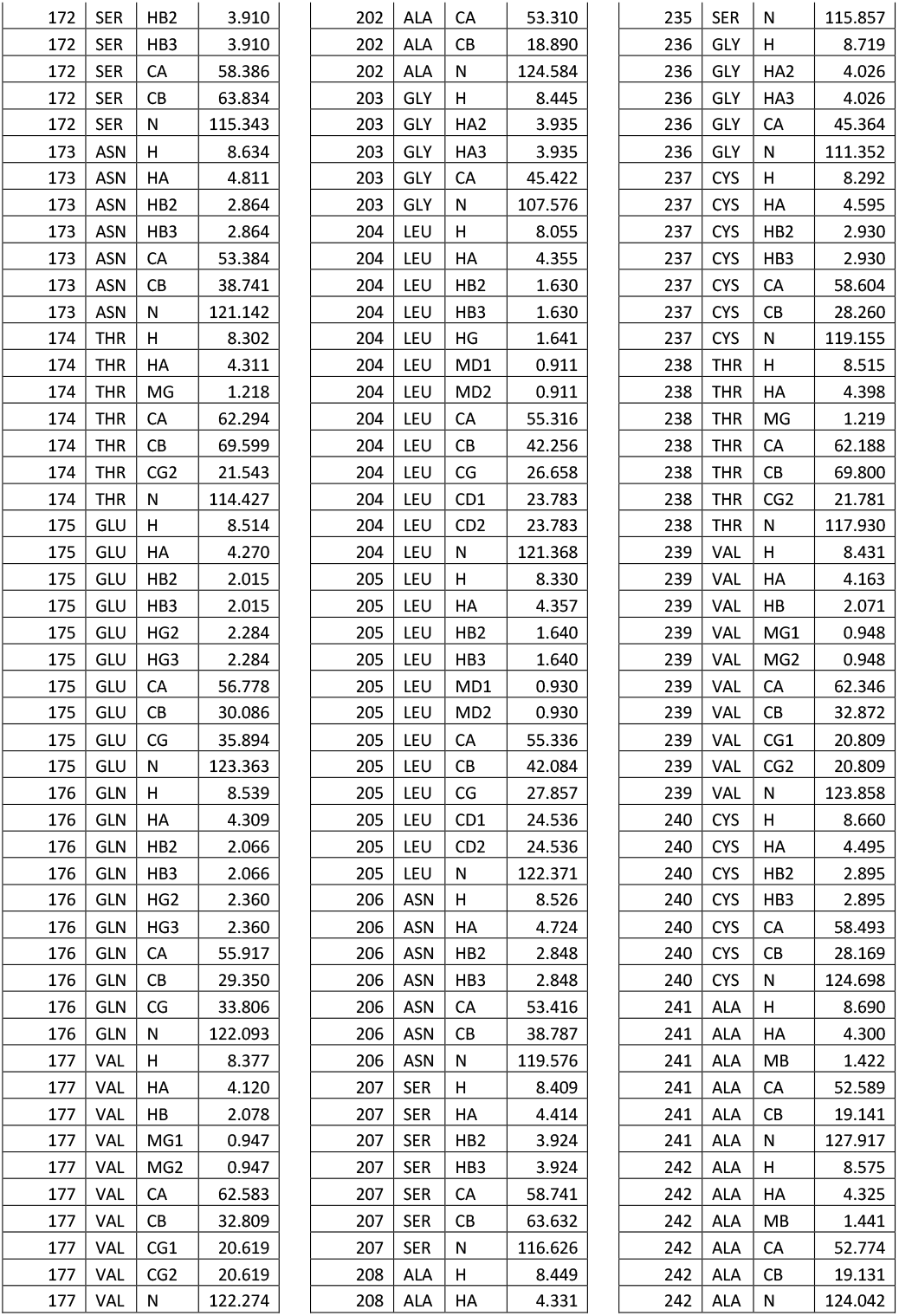
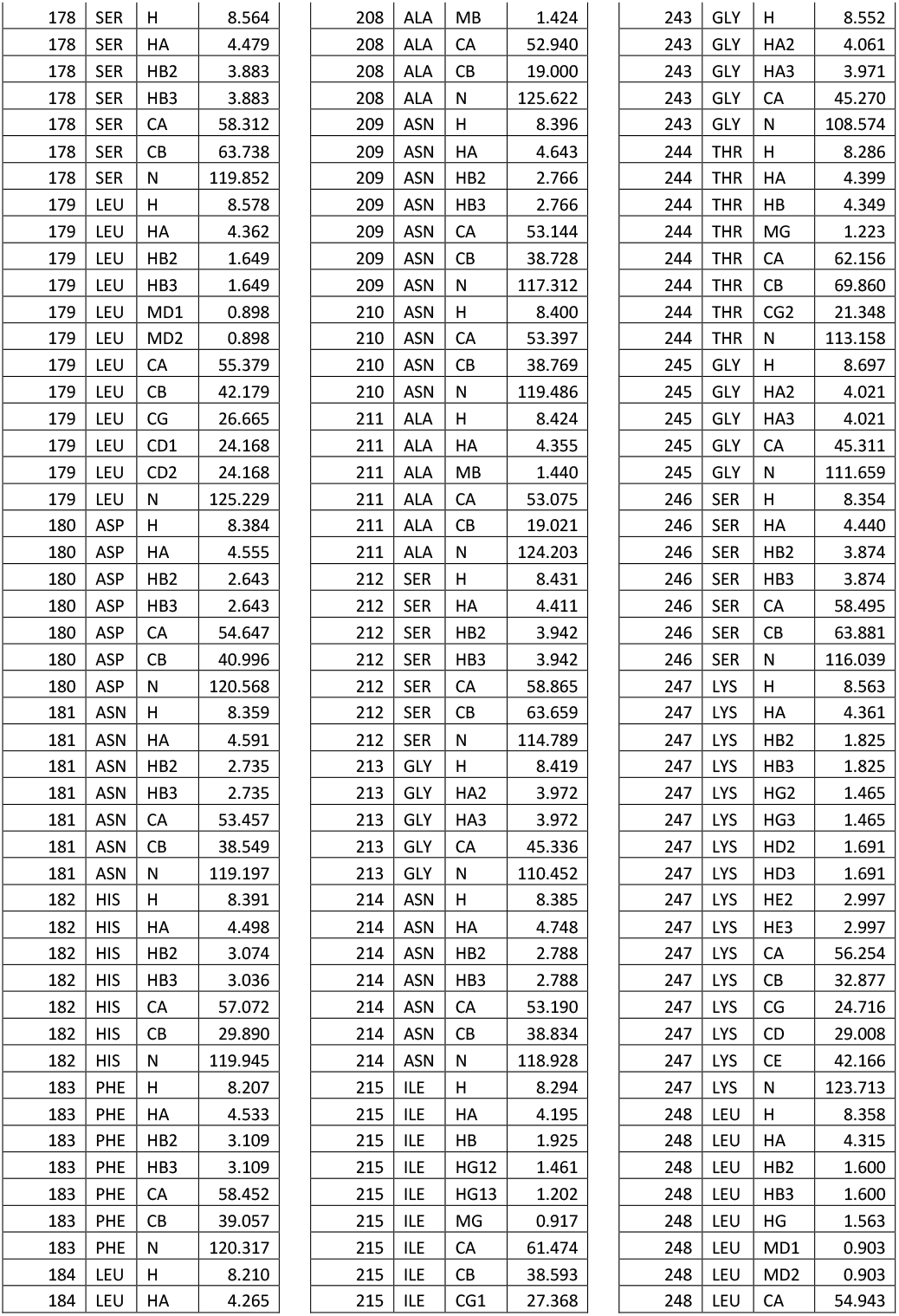
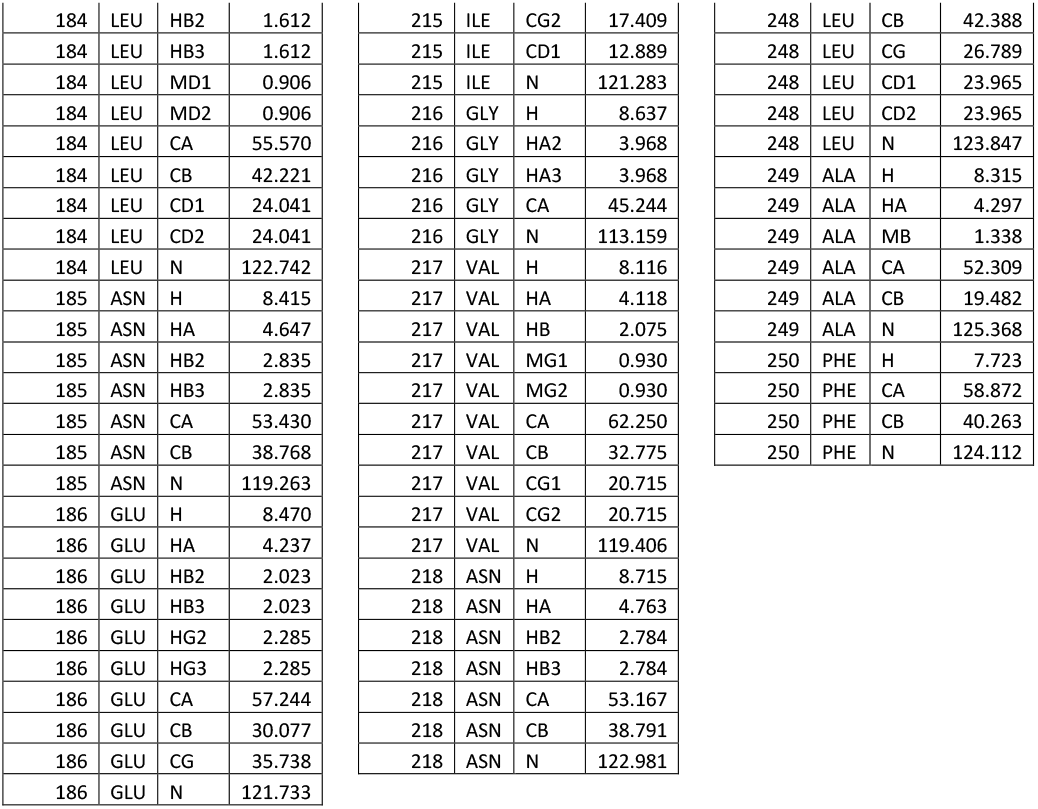
NMR resonance assignments of L2R3C FapC.

| Residue # | AA | Atom | CS (ppm) | Residue # | AA | Atom | CS (ppm) | Residue # | AA | Atom | CS (ppm) |
| --- | --- | --- | --- | --- | --- | --- | --- | --- | --- | --- | --- |
| 157 | ASN | H | 8.687 | 187 | ALA | H | 8.440 | 219 | VAL | H | 8.310 |
| 157 | ASN | HA | 4.728 | 187 | ALA | HA | 4.279 | 219 | VAL | HA | 4.079 |
| 157 | ASN | HB2 | 2.828 | 187 | ALA | MB | 1.432 | 219 | VAL | HB | 2.096 |
| 157 | ASN | HB3 | 2.828 | 187 | ALA | CA | 53.347 | 219 | VAL | MG1 | 0.948 |
| 157 | ASN | CA | 53.294 | 187 | ALA | CB | 18.847 | 219 | VAL | MG2 | 0.948 |
| 157 | ASN | CB | 38.612 | 187 | ALA | N | 124.345 | 219 | VAL | CA | 62.425 |
| 157 | ASN | N | 121.779 | 188 | SER | H | 8.306 | 219 | VAL | CB | 32.663 |
| 158 | LEU | H | 8.423 | 188 | SER | HA | 4.402 | 219 | VAL | CG1 | 20.598 |
| 158 | LEU | HA | 4.331 | 188 | SER | HB2 | 3.924 | 219 | VAL | CG2 | 20.598 |
| 158 | LEU | HB2 | 1.624 | 188 | SER | HB3 | 3.924 | 219 | VAL | N | 121.593 |
| 158 | LEU | HB3 | 1.624 | 188 | SER | CA | 58.789 | 220 | ALA | H | 8.519 |
| 158 | LEU | MD1 | 0.902 | 188 | SER | CB | 63.556 | 220 | ALA | HA | 4.296 |
| 158 | LEU | MD2 | 0.902 | 188 | SER | N | 114.373 | 220 | ALA | MB | 1.407 |
| 158 | LEU | CA | 55.148 | 189 | SER | H | 8.332 | 220 | ALA | CA | 52.584 |
| 158 | LEU | CB | 42.302 | 189 | SER | HA | 4.439 | 220 | ALA | CB | 19.048 |
| 158 | LEU | CG | 26.334 | 189 | SER | HB2 | 3.908 | 220 | ALA | N | 127.867 |
| 158 | LEU | CD1 | 23.673 | 189 | SER | HB3 | 3.908 | 221 | ALA | H | 8.398 |
| 158 | LEU | CD2 | 23.673 | 189 | SER | CA | 58.674 | 221 | ALA | HA | 4.293 |
| 158 | LEU | N | 123.363 | 189 | SER | CB | 63.655 | 221 | ALA | MB | 1.420 |
| 159 | ALA | H | 8.359 | 189 | SER | N | 117.815 | 221 | ALA | CA | 52.793 |
| 159 | ALA | HA | 4.309 | 190 | LYS | H | 8.288 | 221 | ALA | CB | 19.081 |
| 159 | ALA | MB | 1.379 | 190 | LYS | HA | 4.232 | 221 | ALA | N | 123.703 |
| 159 | ALA | CA | 52.402 | 190 | LYS | HB2 | 1.738 | 222 | GLY | H | 8.480 |
| 159 | ALA | CB | 18.997 | 190 | LYS | HB3 | 1.738 | 222 | GLY | HA2 | 3.983 |
| 159 | ALA | N | 124.976 | 190 | LYS | HG2 | 1.358 | 222 | GLY | HA3 | 4.050 |
| 160 | ILE | H | 8.243 | 190 | LYS | HG3 | 1.358 | 222 | GLY | CA | 45.237 |
| 160 | ILE | HA | 4.145 | 190 | LYS | HD2 | 1.695 | 222 | GLY | N | 108.520 |
| 160 | ILE | HB | 1.868 | 190 | LYS | HD3 | 1.695 | 223 | ALA | H | 8.338 |
| 160 | ILE | HG12 | 1.480 | 190 | LYS | HE2 | 2.982 | 223 | ALA | HA | 4.332 |
| 160 | ILE | HG13 | 1.480 | 190 | LYS | HE3 | 2.982 | 223 | ALA | MB | 1.428 |
| 160 | ILE | MD | 0.880 | 190 | LYS | CA | 56.707 | 223 | ALA | CA | 52.786 |
| 160 | ILE | CA | 61.003 | 190 | LYS | CB | 32.625 | 223 | ALA | CB | 19.255 |
| 160 | ILE | CB | 38.634 | 190 | LYS | CG | 24.255 | 223 | ALA | N | 123.852 |
| 160 | ILE | CG1 | 26.598 | 190 | LYS | CD | 29.142 | 224 | GLY | H | 8.594 |
| 160 | ILE | CG2 | 17.136 | 190 | LYS | CE | 42.143 | 224 | GLY | HA2 | 3.965 |
| 160 | ILE | CD1 | 12.670 | 190 | LYS | N | 122.739 | 224 | GLY | HA3 | 3.965 |
| 160 | ILE | N | 121.120 | 191 | HIS | H | 8.276 | 224 | GLY | CA | 45.311 |
| 161 | VAL | H | 8.469 | 191 | HIS | HA | 4.646 | 224 | GLY | N | 108.067 |
| 161 | VAL | HA | 4.304 | 191 | HIS | HB2 | 3.102 | 225 | ASN | H | 8.429 |
| 161 | VAL | HB | 2.070 | 191 | HIS | HB3 | 3.017 | 225 | ASN | HA | 4.741 |
| 161 | VAL | MG1 | 0.944 | 191 | HIS | CA | 56.180 | 225 | ASN | HB2 | 2.836 |
| 161 | VAL | MG2 | 0.944 | 191 | HIS | CB | 30.707 | 225 | ASN | HB3 | 2.836 |
| 161 | VAL | CA | 62.082 | 191 | HIS | N | 119.772 | 225 | ASN | CA | 53.282 |
| 161 | VAL | CB | 32.926 | 192 | SER | H | 8.277 | 225 | ASN | CB | 38.773 |
| 161 | VAL | CG1 | 20.696 | 192 | SER | HA | 4.422 | 225 | ASN | N | 118.699 |
| 161 | VAL | CG2 | 20.696 | 192 | SER | HB2 | 3.821 | 226 | GLN | H | 8.606 |
| 161 | VAL | N | 125.944 | 192 | SER | HB3 | 3.821 | 226 | GLN | HA | 4.332 |
| 162 | THR | H | 8.521 | 192 | SER | CA | 58.361 | 226 | GLN | HB2 | 2.077 |
| 162 | THR | HA | 4.368 | 192 | SER | CB | 63.756 | 226 | GLN | HB3 | 2.077 |
| 162 | THR | HB | 4.272 | 192 | SER | N | 116.875 | 226 | GLN | HG2 | 2.388 |
| 162 | THR | MG | 1.217 | 193 | TYR | H | 8.393 | 226 | GLN | HG3 | 2.388 |
| 162 | THR | CA | 61.605 | 193 | TYR | HA | 4.555 | 226 | GLN | CA | 56.159 |
| 162 | THR | CB | 70.075 | 193 | TYR | HB2 | 2.992 | 226 | GLN | CB | 29.191 |
| 162 | THR | CG2 | 21.632 | 193 | TYR | HB3 | 2.992 | 226 | GLN | CG | 33.699 |
| 162 | THR | N | 119.259 | 193 | TYR | CA | 58.040 | 226 | GLN | N | 120.821 |
| 163 | ALA | H | 8.683 | 193 | TYR | CB | 38.753 | 227 | GLN | H | 8.605 |
| 163 | ALA | HA | 4.316 | 193 | TYR | N | 123.030 | 227 | GLN | HA | 4.359 |
| 163 | ALA | MB | 1.423 | 194 | LYS | H | 8.231 | 227 | GLN | HB2 | 2.094 |
| 163 | ALA | CA | 52.947 | 194 | LYS | CA | 53.622 | 227 | GLN | HB3 | 2.094 |
| 163 | ALA | CB | 19.024 | 194 | LYS | CB | 32.861 | 227 | GLN | HG2 | 2.401 |
| 163 | ALA | N | 126.704 | 194 | LYS | N | 126.241 | 227 | GLN | HG3 | 2.401 |
| 164 | ASP | H | 8.408 | 195 | PRO | HA | 4.285 | 227 | GLN | CA | 56.146 |
| 164 | ASP | HA | 4.584 | 195 | PRO | HB2 | 2.246 | 227 | GLN | CB | 29.340 |
| 164 | ASP | HB2 | 2.738 | 195 | PRO | HB3 | 1.749 | 227 | GLN | CG | 33.881 |
| 164 | ASP | HB3 | 2.738 | 195 | PRO | HG2 | 1.969 | 227 | GLN | N | 121.507 |
| 164 | ASP | CA | 54.291 | 195 | PRO | HG3 | 1.969 | 228 | SER | H | 8.551 |
| 164 | ASP | CB | 40.987 | 195 | PRO | HD2 | 3.575 | 228 | SER | HA | 4.419 |
| 164 | ASP | N | 119.107 | 195 | PRO | HD3 | 3.575 | 228 | SER | HB2 | 3.899 |
| 165 | GLY | H | 8.429 | 195 | PRO | CA | 62.994 | 228 | SER | HB3 | 3.899 |
| 165 | GLY | HA2 | 3.934 | 195 | PRO | CB | 32.031 | 228 | SER | CA | 58.519 |
| 165 | GLY | HA3 | 3.934 | 195 | PRO | CG | 27.313 | 228 | SER | CB | 63.679 |
| 165 | GLY | CA | 45.608 | 195 | PRO | CD | 50.565 | 228 | SER | N | 117.220 |
| 165 | GLY | N | 109.368 | 196 | GLN | H | 8.581 | 229 | ASN | H | 8.663 |
| 166 | LYS | H | 8.211 | 196 | GLN | HA | 4.230 | 229 | ASN | HA | 4.811 |
| 166 | LYS | HA | 4.325 | 196 | GLN | HB2 | 1.952 | 229 | ASN | HB2 | 2.872 |
| 166 | LYS | HB2 | 1.824 | 196 | GLN | HB3 | 1.952 | 229 | ASN | HB3 | 2.872 |
| 166 | LYS | HB3 | 1.824 | 196 | GLN | HG2 | 2.293 | 229 | ASN | CA | 53.361 |

|  |  |  |  |
| --- | --- | --- | --- |
| 166 | LYS | HG2 | 1.397 |
| 166 | LYS | HG3 | 1.397 |
| 166 | LYS | HD2 | 1.700 |
| 166 | LYS | HD3 | 1.700 |
| 166 | LYS | HE2 | 2.990 |
| 166 | LYS | HE3 | 2.990 |
| 166 | LYS | CA | 56.343 |
| 166 | LYS | CB | 32.835 |
| 166 | LYS | CG | 25.000 |
| 166 | LYS | CD | 29.014 |
| 166 | LYS | CE | 42.066 |
| 166 | LYS | N | 120.474 |
| 167 | ASN | H | 8.616 |
| 167 | ASN | HA | 4.753 |
| 167 | ASN | HB2 | 2.792 |
| 167 | ASN | HB3 | 2.792 |
| 167 | ASN | CA | 53.362 |
| 167 | ASN | CB | 38.769 |
| 167 | ASN | N | 119.764 |
| 168 | VAL | H | 8.270 |
| 168 | VAL | HA | 4.085 |
| 168 | VAL | HB | 2.084 |
| 168 | VAL | MG1 | 0.943 |
| 168 | VAL | MG2 | 0.943 |
| 168 | VAL | CA | 62.354 |
| 168 | VAL | CB | 32.694 |
| 168 | VAL | CG1 | 20.628 |
| 168 | VAL | CG2 | 20.628 |
| 168 | VAL | N | 121.320 |
| 169 | ALA | H | 8.534 |
| 169 | ALA | HA | 4.301 |
| 169 | ALA | MB | 1.404 |
| 169 | ALA | CA | 52.440 |
| 169 | ALA | CB | 19.080 |
| 169 | ALA | N | 128.120 |
| 170 | ALA | H | 8.409 |
| 170 | ALA | HA | 4.275 |
| 170 | ALA | MB | 1.413 |
| 170 | ALA | CA | 52.363 |
| 170 | ALA | CB | 19.150 |
| 170 | ALA | N | 123.908 |
| 171 | ALA | H | 8.474 |
| 171 | ALA | HA | 4.331 |
| 171 | ALA | MB | 1.428 |
| 171 | ALA | CA | 52.530 |
| 171 | ALA | CB | 19.189 |
| 171 | ALA | N | 123.879 |
| 172 | SER | H | 8.470 |
| 172 | SER | HA | 4.449 |
| 196 | GLN | HG3 | 2.293 |
| 196 | GLN | CA | 55.768 |
| 196 | GLN | CB | 29.569 |
| 196 | GLN | CG | 33.608 |
| 196 | GLN | N | 120.566 |
| 197 | TYR | H | 8.297 |
| 197 | TYR | HA | 4.638 |
| 197 | TYR | HB2 | 2.969 |
| 197 | TYR | HB3 | 2.969 |
| 197 | TYR | CA | 57.534 |
| 197 | TYR | CB | 38.972 |
| 197 | TYR | N | 121.602 |
| 198 | VAL | H | 8.217 |
| 198 | VAL | HA | 4.074 |
| 198 | VAL | HB | 1.969 |
| 198 | VAL | MG1 | 0.898 |
| 198 | VAL | MG2 | 0.898 |
| 198 | VAL | CA | 62.054 |
| 198 | VAL | CB | 33.099 |
| 198 | VAL | CG1 | 20.868 |
| 198 | VAL | CG2 | 20.868 |
| 198 | VAL | N | 123.829 |
| 199 | VAL | H | 8.388 |
| 199 | VAL | HA | 4.034 |
| 199 | VAL | HB | 2.052 |
| 199 | VAL | MG1 | 0.946 |
| 199 | VAL | MG2 | 0.946 |
| 199 | VAL | CA | 62.352 |
| 199 | VAL | CB | 32.710 |
| 199 | VAL | CG1 | 20.847 |
| 199 | VAL | CG2 | 20.847 |
| 199 | VAL | N | 125.296 |
| 200 | ASN | H | 8.695 |
| 200 | ASN | HA | 4.741 |
| 200 | ASN | HB2 | 2.842 |
| 200 | ASN | HB3 | 2.842 |
| 200 | ASN | CA | 52.932 |
| 200 | ASN | CB | 38.925 |
| 200 | ASN | N | 123.453 |
| 201 | ASN | H | 8.630 |
| 201 | ASN | HA | 4.707 |
| 201 | ASN | HB2 | 2.823 |
| 201 | ASN | HB3 | 2.823 |
| 201 | ASN | CA | 53.191 |
| 201 | ASN | CB | 38.816 |
| 201 | ASN | N | 120.662 |
| 202 | ALA | H | 8.438 |
| 202 | ALA | HA | 4.272 |
| 202 | ALA | MB | 1.428 |
| 229 | ASN | CB | 38.724 |
| 229 | ASN | N | 121.273 |
| 230 | THR | H | 8.278 |
| 230 | THR | HA | 4.288 |
| 230 | THR | MG | 1.205 |
| 230 | THR | CA | 62.405 |
| 230 | THR | CB | 69.627 |
| 230 | THR | CG2 | 21.340 |
| 230 | THR | N | 114.763 |
| 231 | LEU | H | 8.396 |
| 231 | LEU | HA | 4.434 |
| 231 | LEU | HB2 | 1.654 |
| 231 | LEU | HB3 | 1.654 |
| 231 | LEU | MD1 | 0.922 |
| 231 | LEU | MD2 | 0.922 |
| 231 | LEU | CA | 55.401 |
| 231 | LEU | CB | 42.257 |
| 231 | LEU | CG | 26.803 |
| 231 | LEU | CD1 | 23.935 |
| 231 | LEU | CD2 | 23.935 |
| 231 | LEU | N | 124.645 |
| 232 | THR | H | 8.297 |
| 232 | THR | HA | 4.331 |
| 232 | THR | HB | 4.251 |
| 232 | THR | MG | 1.207 |
| 232 | THR | CA | 61.971 |
| 232 | THR | CB | 69.734 |
| 232 | THR | CG2 | 21.316 |
| 232 | THR | N | 116.074 |
| 233 | LEU | H | 8.469 |
| 233 | LEU | HA | 4.371 |
| 233 | LEU | HB2 | 1.655 |
| 233 | LEU | HB3 | 1.655 |
| 233 | LEU | MD1 | 0.914 |
| 233 | LEU | MD2 | 0.914 |
| 233 | LEU | CA | 55.372 |
| 233 | LEU | CB | 42.317 |
| 233 | LEU | CG | 26.371 |
| 233 | LEU | CD1 | 23.676 |
| 233 | LEU | CD2 | 23.676 |
| 233 | LEU | N | 125.492 |
| 234 | GLY | H | 8.615 |
| 234 | GLY | HA2 | 4.017 |
| 234 | GLY | HA3 | 4.017 |
| 234 | GLY | CA | 45.264 |
| 234 | GLY | N | 110.287 |
| 235 | SER | H | 8.452 |
| 235 | SER | CA | 58.696 |
| 235 | SER | CB | 63.814 |
| 172 | SER | HB2 | 3.910 |
| 172 | SER | HB3 | 3.910 |
| 172 | SER | CA | 58.386 |
| 172 | SER | CB | 63.834 |
| 172 | SER | N | 115.343 |
| 173 | ASN | H | 8.634 |
| 173 | ASN | HA | 4.811 |
| 173 | ASN | HB2 | 2.864 |
| 173 | ASN | HB3 | 2.864 |
| 173 | ASN | CA | 53.384 |
| 173 | ASN | CB | 38.741 |
| 173 | ASN | N | 121.142 |
| 174 | THR | H | 8.302 |
| 174 | THR | HA | 4.311 |
| 174 | THR | MG | 1.218 |
| 174 | THR | CA | 62.294 |
| 174 | THR | CB | 69.599 |
| 174 | THR | CG2 | 21.543 |
| 174 | THR | N | 114.427 |
| 175 | GLU | H | 8.514 |
| 175 | GLU | HA | 4.270 |
| 175 | GLU | HB2 | 2.015 |
| 175 | GLU | HB3 | 2.015 |
| 175 | GLU | HG2 | 2.284 |
| 175 | GLU | HG3 | 2.284 |
| 175 | GLU | CA | 56.778 |
| 175 | GLU | CB | 30.086 |
| 175 | GLU | CG | 35.894 |
| 175 | GLU | N | 123.363 |
| 176 | GLN | H | 8.539 |
| 176 | GLN | HA | 4.309 |
| 176 | GLN | HB2 | 2.066 |
| 176 | GLN | HB3 | 2.066 |
| 176 | GLN | HG2 | 2.360 |
| 176 | GLN | HG3 | 2.360 |
| 176 | GLN | CA | 55.917 |
| 176 | GLN | CB | 29.350 |
| 176 | GLN | CG | 33.806 |
| 176 | GLN | N | 122.093 |
| 177 | VAL | H | 8.377 |
| 177 | VAL | HA | 4.120 |
| 177 | VAL | HB | 2.078 |
| 177 | VAL | MG1 | 0.947 |
| 177 | VAL | MG2 | 0.947 |
| 177 | VAL | CA | 62.583 |
| 177 | VAL | CB | 32.809 |
| 177 | VAL | CG1 | 20.619 |
| 177 | VAL | CG2 | 20.619 |
| 177 | VAL | N | 122.274 |
| 202 | ALA | CA | 53.310 |
| 202 | ALA | CB | 18.890 |
| 202 | ALA | N | 124.584 |
| 203 | GLY | H | 8.445 |
| 203 | GLY | HA2 | 3.935 |
| 203 | GLY | HA3 | 3.935 |
| 203 | GLY | CA | 45.422 |
| 203 | GLY | N | 107.576 |
| 204 | LEU | H | 8.055 |
| 204 | LEU | HA | 4.355 |
| 204 | LEU | HB2 | 1.630 |
| 204 | LEU | HB3 | 1.630 |
| 204 | LEU | HG | 1.641 |
| 204 | LEU | MD1 | 0.911 |
| 204 | LEU | MD2 | 0.911 |
| 204 | LEU | CA | 55.316 |
| 204 | LEU | CB | 42.256 |
| 204 | LEU | CG | 26.658 |
| 204 | LEU | CD1 | 23.783 |
| 204 | LEU | CD2 | 23.783 |
| 204 | LEU | N | 121.368 |
| 205 | LEU | H | 8.330 |
| 205 | LEU | HA | 4.357 |
| 205 | LEU | HB2 | 1.640 |
| 205 | LEU | HB3 | 1.640 |
| 205 | LEU | MD1 | 0.930 |
| 205 | LEU | MD2 | 0.930 |
| 205 | LEU | CA | 55.336 |
| 205 | LEU | CB | 42.084 |
| 205 | LEU | CG | 27.857 |
| 205 | LEU | CD1 | 24.536 |
| 205 | LEU | CD2 | 24.536 |
| 205 | LEU | N | 122.371 |
| 206 | ASN | H | 8.526 |
| 206 | ASN | HA | 4.724 |
| 206 | ASN | HB2 | 2.848 |
| 206 | ASN | HB3 | 2.848 |
| 206 | ASN | CA | 53.416 |
| 206 | ASN | CB | 38.787 |
| 206 | ASN | N | 119.576 |
| 207 | SER | H | 8.409 |
| 207 | SER | HA | 4.414 |
| 207 | SER | HB2 | 3.924 |
| 207 | SER | HB3 | 3.924 |
| 207 | SER | CA | 58.741 |
| 207 | SER | CB | 63.632 |
| 207 | SER | N | 116.626 |
| 208 | ALA | H | 8.449 |
| 208 | ALA | HA | 4.331 |
| 235 | SER | N | 115.857 |
| 236 | GLY | H | 8.719 |
| 236 | GLY | HA2 | 4.026 |
| 236 | GLY | HA3 | 4.026 |
| 236 | GLY | CA | 45.364 |
| 236 | GLY | N | 111.352 |
| 237 | CYS | H | 8.292 |
| 237 | CYS | HA | 4.595 |
| 237 | CYS | HB2 | 2.930 |
| 237 | CYS | HB3 | 2.930 |
| 237 | CYS | CA | 58.604 |
| 237 | CYS | CB | 28.260 |
| 237 | CYS | N | 119.155 |
| 238 | THR | H | 8.515 |
| 238 | THR | HA | 4.398 |
| 238 | THR | MG | 1.219 |
| 238 | THR | CA | 62.188 |
| 238 | THR | CB | 69.800 |
| 238 | THR | CG2 | 21.781 |
| 238 | THR | N | 117.930 |
| 239 | VAL | H | 8.431 |
| 239 | VAL | HA | 4.163 |
| 239 | VAL | HB | 2.071 |
| 239 | VAL | MG1 | 0.948 |
| 239 | VAL | MG2 | 0.948 |
| 239 | VAL | CA | 62.346 |
| 239 | VAL | CB | 32.872 |
| 239 | VAL | CG1 | 20.809 |
| 239 | VAL | CG2 | 20.809 |
| 239 | VAL | N | 123.858 |
| 240 | CYS | H | 8.660 |
| 240 | CYS | HA | 4.495 |
| 240 | CYS | HB2 | 2.895 |
| 240 | CYS | HB3 | 2.895 |
| 240 | CYS | CA | 58.493 |
| 240 | CYS | CB | 28.169 |
| 240 | CYS | N | 124.698 |
| 241 | ALA | H | 8.690 |
| 241 | ALA | HA | 4.300 |
| 241 | ALA | MB | 1.422 |
| 241 | ALA | CA | 52.589 |
| 241 | ALA | CB | 19.141 |
| 241 | ALA | N | 127.917 |
| 242 | ALA | H | 8.575 |
| 242 | ALA | HA | 4.325 |
| 242 | ALA | MB | 1.441 |
| 242 | ALA | CA | 52.774 |
| 242 | ALA | CB | 19.131 |
| 242 | ALA | N | 124.042 |
| 178 | SER | H | 8.564 |
| 178 | SER | HA | 4.479 |
| 178 | SER | HB2 | 3.883 |
| 178 | SER | HB3 | 3.883 |
| 178 | SER | CA | 58.312 |
| 178 | SER | CB | 63.738 |
| 178 | SER | N | 119.852 |
| 179 | LEU | H | 8.578 |
| 179 | LEU | HA | 4.362 |
| 179 | LEU | HB2 | 1.649 |
| 179 | LEU | HB3 | 1.649 |
| 179 | LEU | MD1 | 0.898 |
| 179 | LEU | MD2 | 0.898 |
| 179 | LEU | CA | 55.379 |
| 179 | LEU | CB | 42.179 |
| 179 | LEU | CG | 26.665 |
| 179 | LEU | CD1 | 24.168 |
| 179 | LEU | CD2 | 24.168 |
| 179 | LEU | N | 125.229 |
| 180 | ASP | H | 8.384 |
| 180 | ASP | HA | 4.555 |
| 180 | ASP | HB2 | 2.643 |
| 180 | ASP | HB3 | 2.643 |
| 180 | ASP | CA | 54.647 |
| 180 | ASP | CB | 40.996 |
| 180 | ASP | N | 120.568 |
| 181 | ASN | H | 8.359 |
| 181 | ASN | HA | 4.591 |
| 181 | ASN | HB2 | 2.735 |
| 181 | ASN | HB3 | 2.735 |
| 181 | ASN | CA | 53.457 |
| 181 | ASN | CB | 38.549 |
| 181 | ASN | N | 119.197 |
| 182 | HIS | H | 8.391 |
| 182 | HIS | HA | 4.498 |
| 182 | HIS | HB2 | 3.074 |
| 182 | HIS | HB3 | 3.036 |
| 182 | HIS | CA | 57.072 |
| 182 | HIS | CB | 29.890 |
| 182 | HIS | N | 119.945 |
| 183 | PHE | H | 8.207 |
| 183 | PHE | HA | 4.533 |
| 183 | PHE | HB2 | 3.109 |
| 183 | PHE | HB3 | 3.109 |
| 183 | PHE | CA | 58.452 |
| 183 | PHE | CB | 39.057 |
| 183 | PHE | N | 120.317 |
| 184 | LEU | H | 8.210 |
| 184 | LEU | HA | 4.265 |
| 208 | ALA | MB | 1.424 |
| 208 | ALA | CA | 52.940 |
| 208 | ALA | CB | 19.000 |
| 208 | ALA | N | 125.622 |
| 209 | ASN | H | 8.396 |
| 209 | ASN | HA | 4.643 |
| 209 | ASN | HB2 | 2.766 |
| 209 | ASN | HB3 | 2.766 |
| 209 | ASN | CA | 53.144 |
| 209 | ASN | CB | 38.728 |
| 209 | ASN | N | 117.312 |
| 210 | ASN | H | 8.400 |
| 210 | ASN | CA | 53.397 |
| 210 | ASN | CB | 38.769 |
| 210 | ASN | N | 119.486 |
| 211 | ALA | H | 8.424 |
| 211 | ALA | HA | 4.355 |
| 211 | ALA | MB | 1.440 |
| 211 | ALA | CA | 53.075 |
| 211 | ALA | CB | 19.021 |
| 211 | ALA | N | 124.203 |
| 212 | SER | H | 8.431 |
| 212 | SER | HA | 4.411 |
| 212 | SER | HB2 | 3.942 |
| 212 | SER | HB3 | 3.942 |
| 212 | SER | CA | 58.865 |
| 212 | SER | CB | 63.659 |
| 212 | SER | N | 114.789 |
| 213 | GLY | H | 8.419 |
| 213 | GLY | HA2 | 3.972 |
| 213 | GLY | HA3 | 3.972 |
| 213 | GLY | CA | 45.336 |
| 213 | GLY | N | 110.452 |
| 214 | ASN | H | 8.385 |
| 214 | ASN | HA | 4.748 |
| 214 | ASN | HB2 | 2.788 |
| 214 | ASN | HB3 | 2.788 |
| 214 | ASN | CA | 53.190 |
| 214 | ASN | CB | 38.834 |
| 214 | ASN | N | 118.928 |
| 215 | ILE | H | 8.294 |
| 215 | ILE | HA | 4.195 |
| 215 | ILE | HB | 1.925 |
| 215 | ILE | HG12 | 1.461 |
| 215 | ILE | HG13 | 1.202 |
| 215 | ILE | MG | 0.917 |
| 215 | ILE | CA | 61.474 |
| 215 | ILE | CB | 38.593 |
| 215 | ILE | CG1 | 27.368 |
| 243 | GLY | H | 8.552 |
| 243 | GLY | HA2 | 4.061 |
| 243 | GLY | HA3 | 3.971 |
| 243 | GLY | CA | 45.270 |
| 243 | GLY | N | 108.574 |
| 244 | THR | H | 8.286 |
| 244 | THR | HA | 4.399 |
| 244 | THR | HB | 4.349 |
| 244 | THR | MG | 1.223 |
| 244 | THR | CA | 62.156 |
| 244 | THR | CB | 69.860 |
| 244 | THR | CG2 | 21.348 |
| 244 | THR | N | 113.158 |
| 245 | GLY | H | 8.697 |
| 245 | GLY | HA2 | 4.021 |
| 245 | GLY | HA3 | 4.021 |
| 245 | GLY | CA | 45.311 |
| 245 | GLY | N | 111.659 |
| 246 | SER | H | 8.354 |
| 246 | SER | HA | 4.440 |
| 246 | SER | HB2 | 3.874 |
| 246 | SER | HB3 | 3.874 |
| 246 | SER | CA | 58.495 |
| 246 | SER | CB | 63.881 |
| 246 | SER | N | 116.039 |
| 247 | LYS | H | 8.563 |
| 247 | LYS | HA | 4.361 |
| 247 | LYS | HB2 | 1.825 |
| 247 | LYS | HB3 | 1.825 |
| 247 | LYS | HG2 | 1.465 |
| 247 | LYS | HG3 | 1.465 |
| 247 | LYS | HD2 | 1.691 |
| 247 | LYS | HD3 | 1.691 |
| 247 | LYS | HE2 | 2.997 |
| 247 | LYS | HE3 | 2.997 |
| 247 | LYS | CA | 56.254 |
| 247 | LYS | CB | 32.877 |
| 247 | LYS | CG | 24.716 |
| 247 | LYS | CD | 29.008 |
| 247 | LYS | CE | 42.166 |
| 247 | LYS | N | 123.713 |
| 248 | LEU | H | 8.358 |
| 248 | LEU | HA | 4.315 |
| 248 | LEU | HB2 | 1.600 |
| 248 | LEU | HB3 | 1.600 |
| 248 | LEU | HG | 1.563 |
| 248 | LEU | MD1 | 0.903 |
| 248 | LEU | MD2 | 0.903 |
| 248 | LEU | CA | 54.943 |
| 184 | LEU | HB2 | 1.612 |
| 184 | LEU | HB3 | 1.612 |
| 184 | LEU | MD1 | 0.906 |
| 184 | LEU | MD2 | 0.906 |
| 184 | LEU | CA | 55.570 |
| 184 | LEU | CB | 42.221 |
| 184 | LEU | CD1 | 24.041 |
| 184 | LEU | CD2 | 24.041 |
| 184 | LEU | N | 122.742 |
| 185 | ASN | H | 8.415 |
| 185 | ASN | HA | 4.647 |
| 185 | ASN | HB2 | 2.835 |
| 185 | ASN | HB3 | 2.835 |
| 185 | ASN | CA | 53.430 |
| 185 | ASN | CB | 38.768 |
| 185 | ASN | N | 119.263 |
| 186 | GLU | H | 8.470 |
| 186 | GLU | HA | 4.237 |
| 186 | GLU | HB2 | 2.023 |
| 186 | GLU | HB3 | 2.023 |
| 186 | GLU | HG2 | 2.285 |
| 186 | GLU | HG3 | 2.285 |
| 186 | GLU | CA | 57.244 |
| 186 | GLU | CB | 30.077 |
| 186 | GLU | CG | 35.738 |
| 186 | GLU | N | 121.733 |
| 215 | ILE | CG2 | 17.409 |
| 215 | ILE | CD1 | 12.889 |
| 215 | ILE | N | 121.283 |
| 216 | GLY | H | 8.637 |
| 216 | GLY | HA2 | 3.968 |
| 216 | GLY | HA3 | 3.968 |
| 216 | GLY | CA | 45.244 |
| 216 | GLY | N | 113.159 |
| 217 | VAL | H | 8.116 |
| 217 | VAL | HA | 4.118 |
| 217 | VAL | HB | 2.075 |
| 217 | VAL | MG1 | 0.930 |
| 217 | VAL | MG2 | 0.930 |
| 217 | VAL | CA | 62.250 |
| 217 | VAL | CB | 32.775 |
| 217 | VAL | CG1 | 20.715 |
| 217 | VAL | CG2 | 20.715 |
| 217 | VAL | N | 119.406 |
| 218 | ASN | H | 8.715 |
| 218 | ASN | HA | 4.763 |
| 218 | ASN | HB2 | 2.784 |
| 218 | ASN | HB3 | 2.784 |
| 218 | ASN | CA | 53.167 |
| 218 | ASN | CB | 38.791 |
| 218 | ASN | N | 122.981 |
| 248 | LEU | CB | 42.388 |
| 248 | LEU | CG | 26.789 |
| 248 | LEU | CD1 | 23.965 |
| 248 | LEU | CD2 | 23.965 |
| 248 | LEU | N | 123.847 |
| 249 | ALA | H | 8.315 |
| 249 | ALA | HA | 4.297 |
| 249 | ALA | MB | 1.338 |
| 249 | ALA | CA | 52.309 |
| 249 | ALA | CB | 19.482 |
| 249 | ALA | N | 125.368 |
| 250 | PHE | H | 7.723 |
| 250 | PHE | CA | 58.872 |
| 250 | PHE | CB | 40.263 |
| 250 | PHE | N | 124.112 |

**Table 2.**
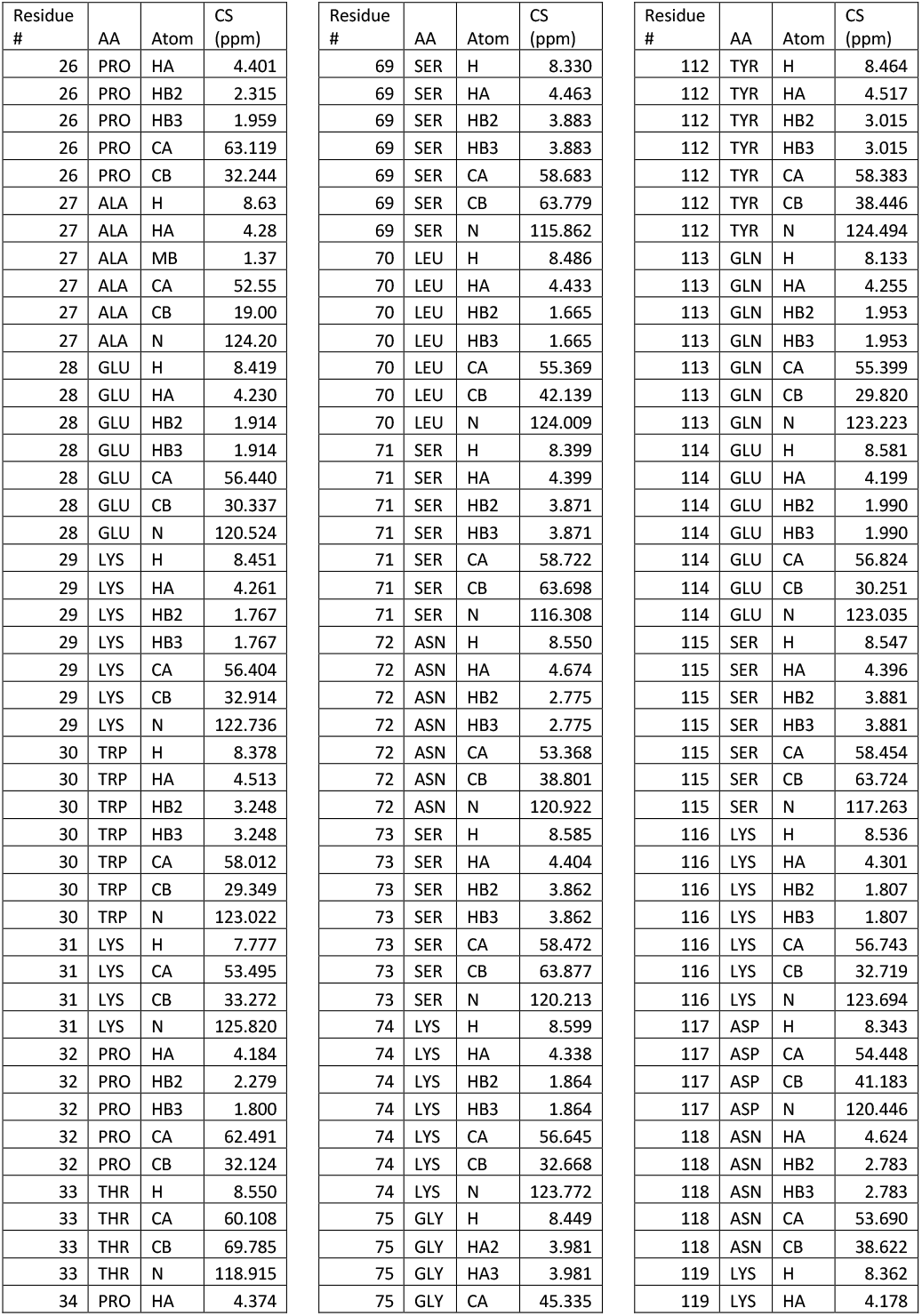
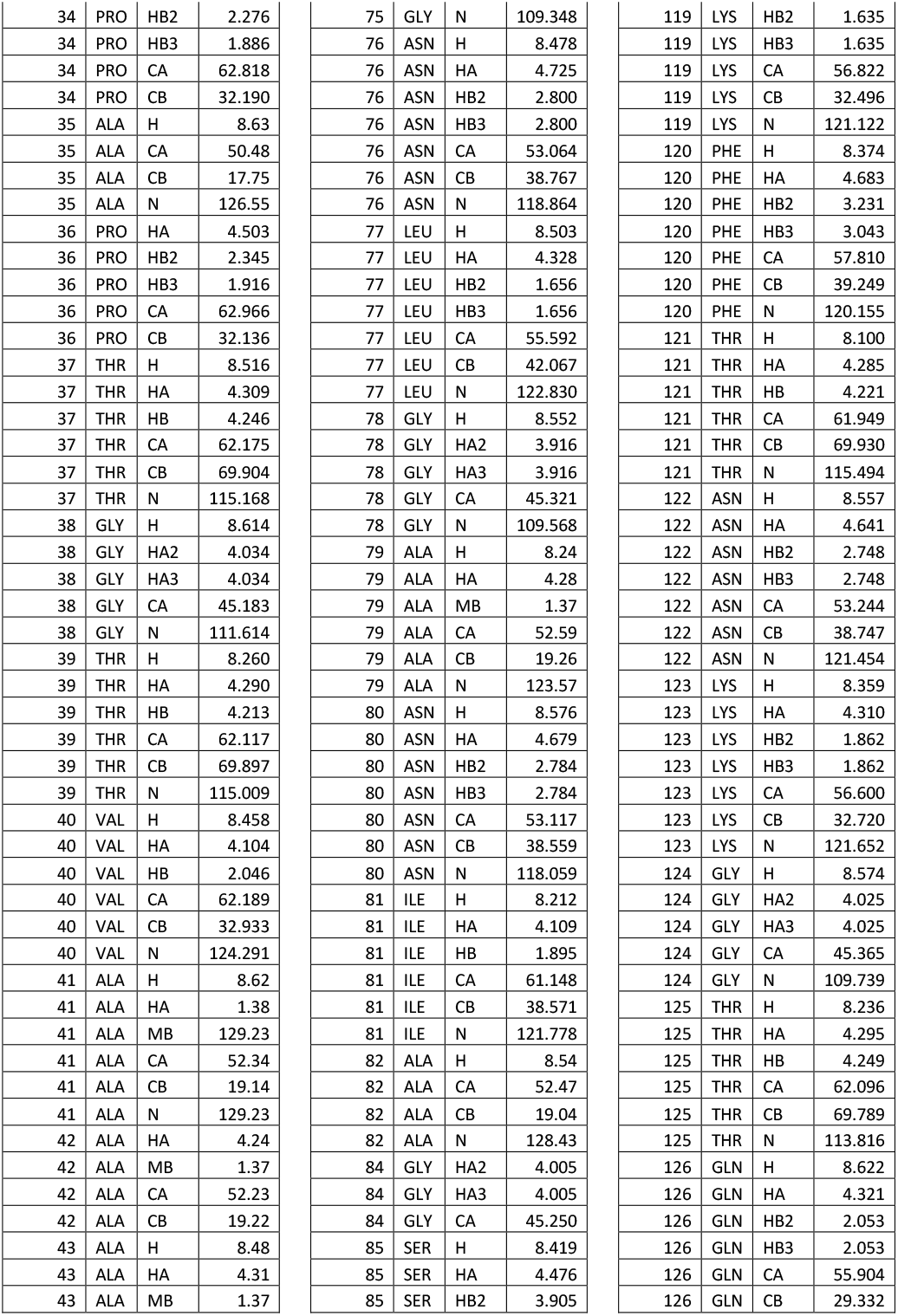
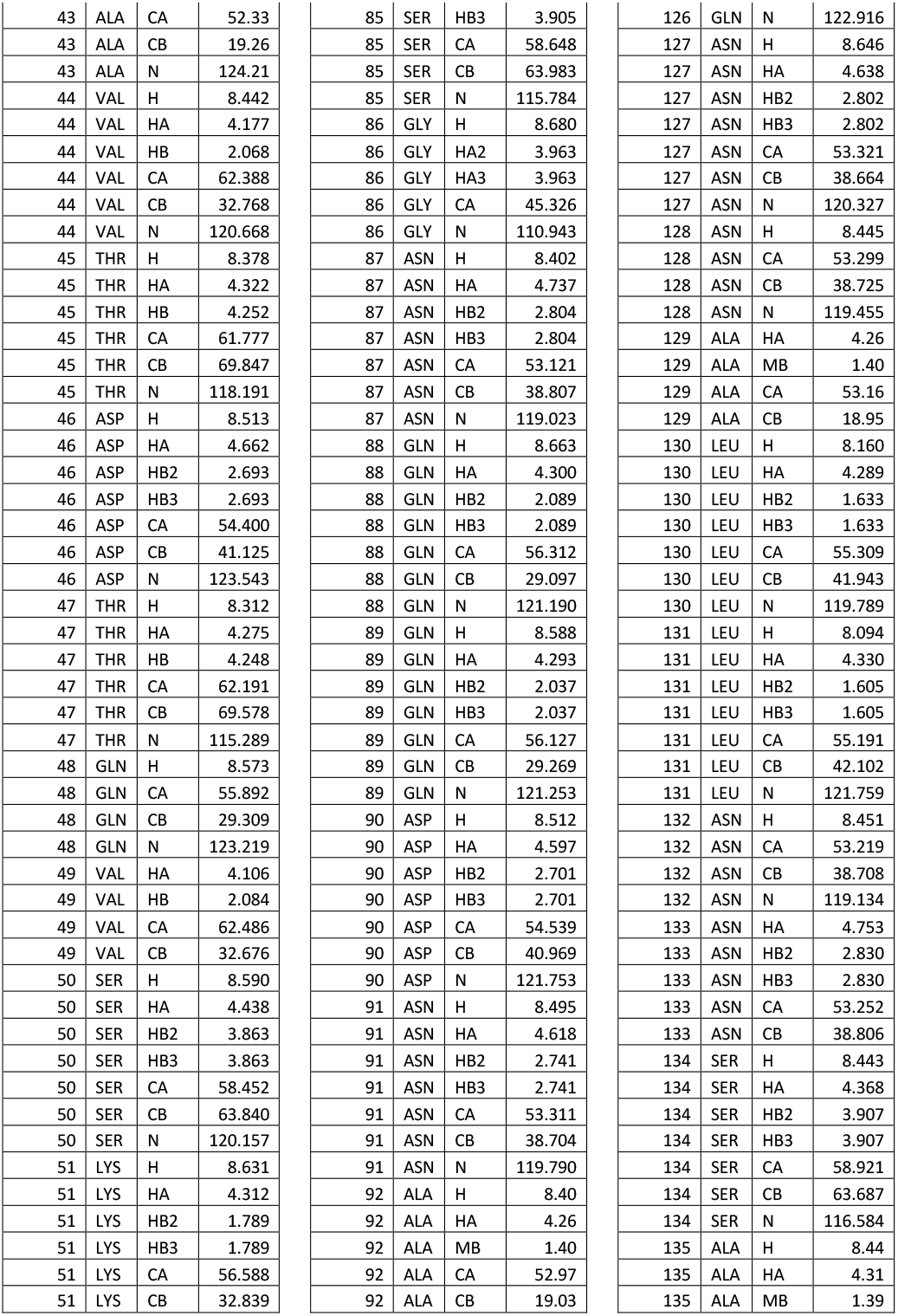
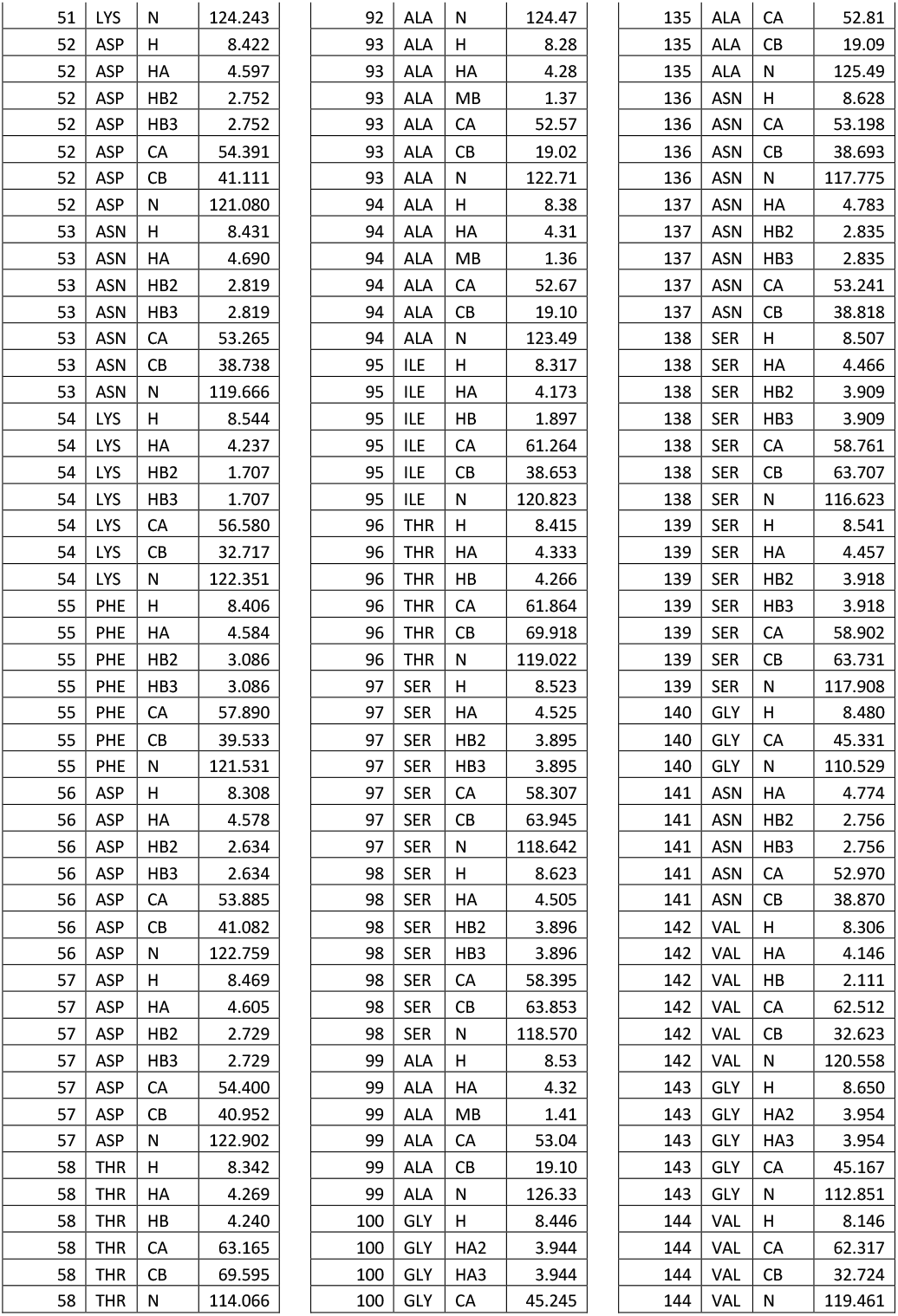
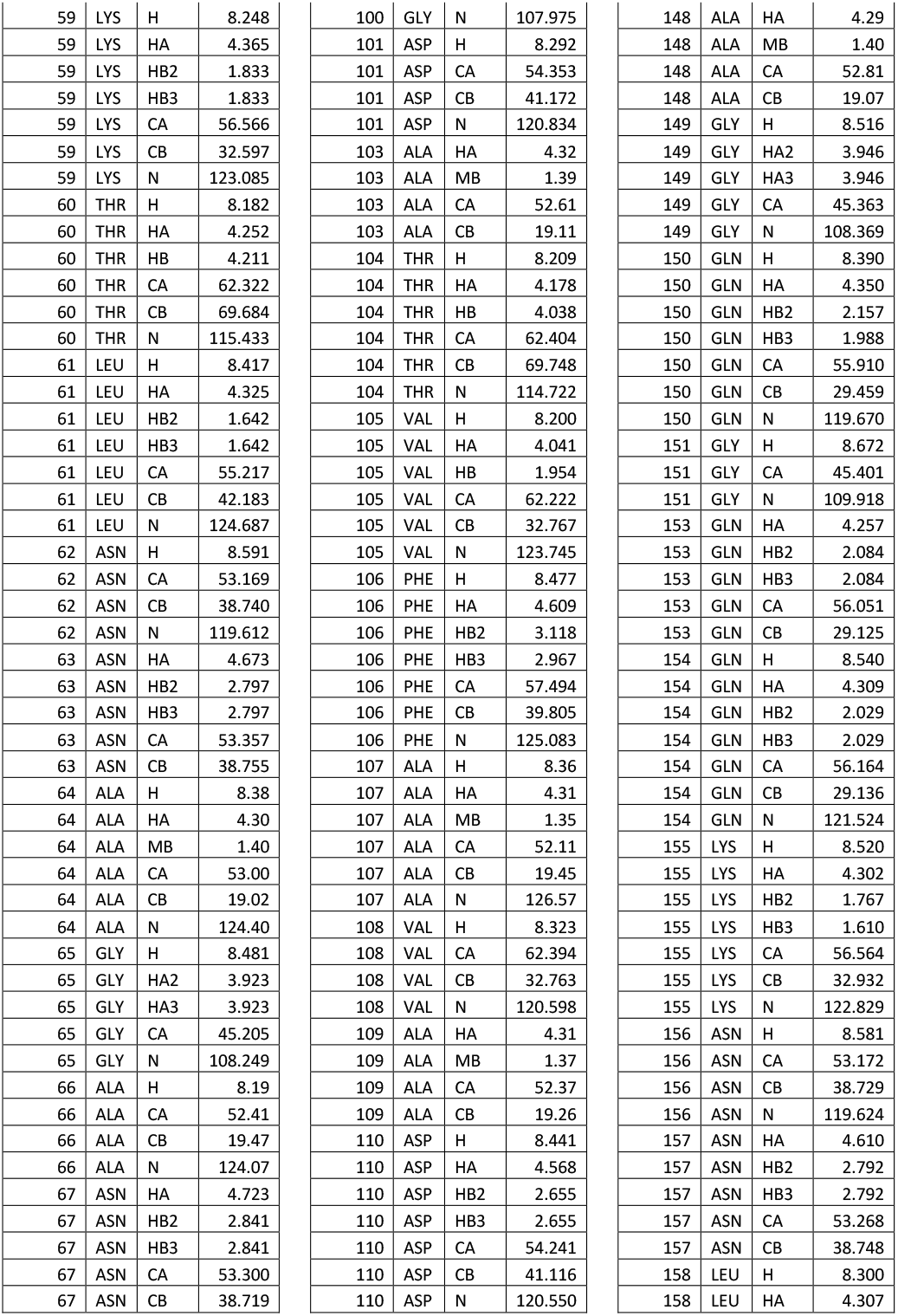
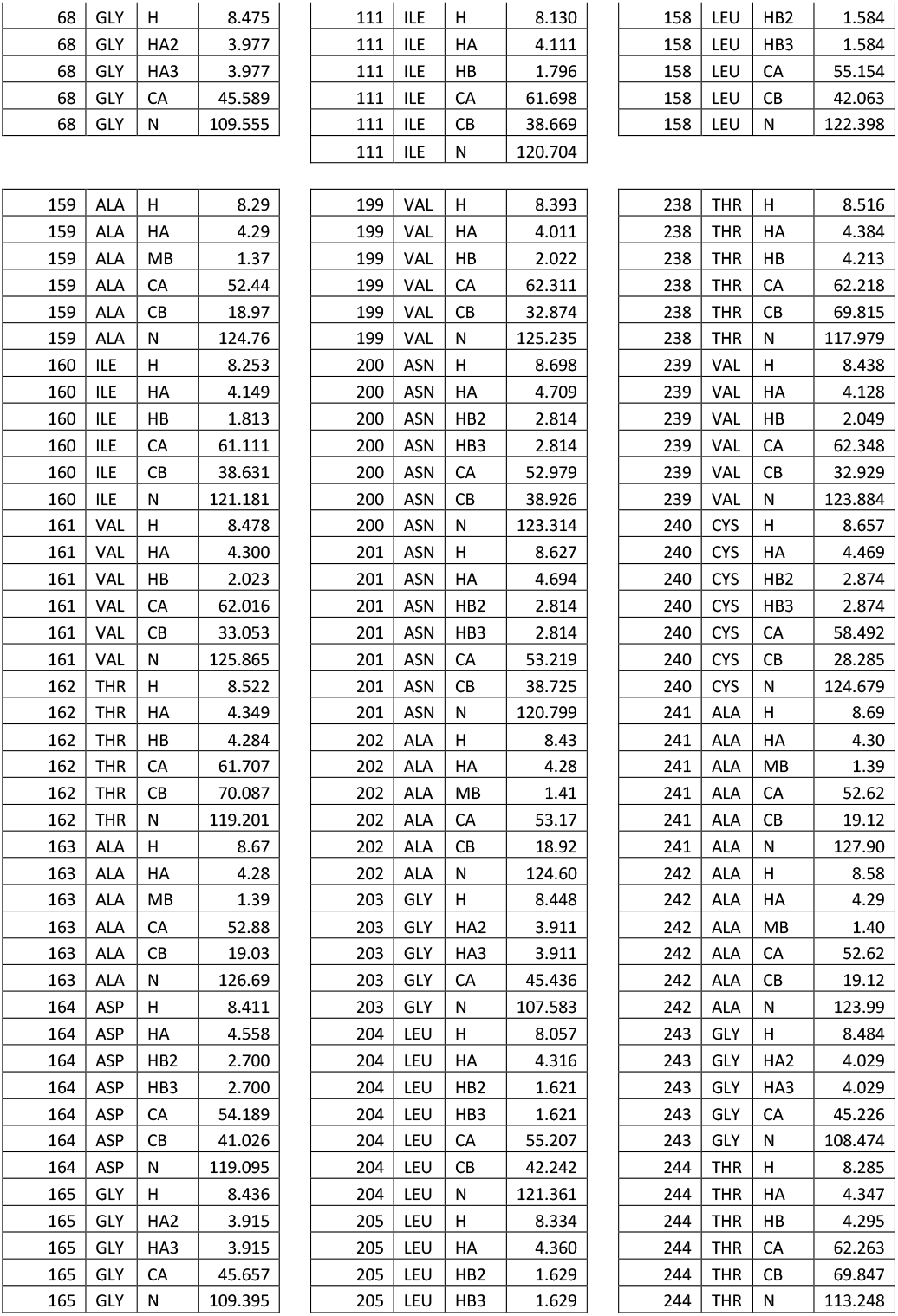
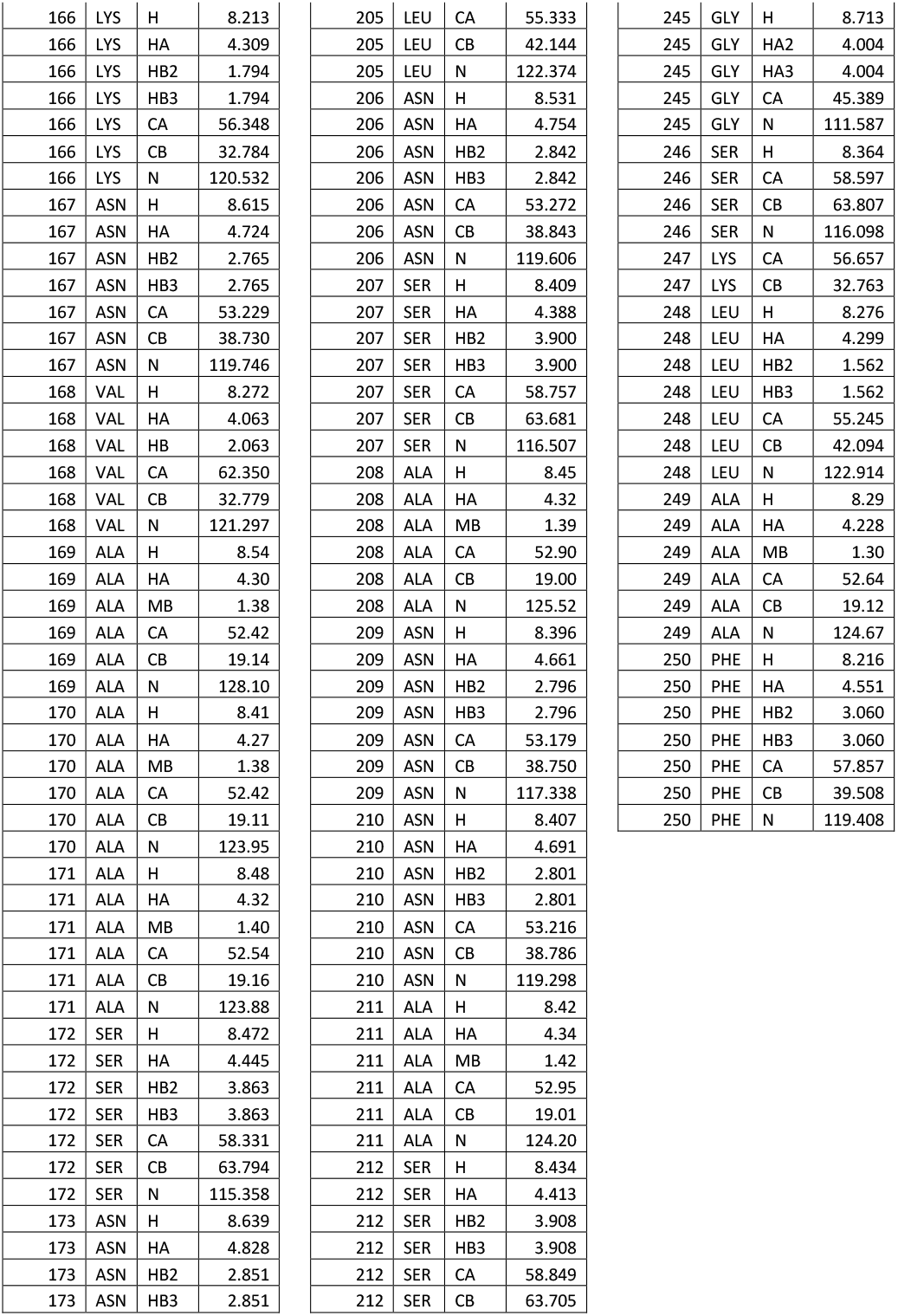
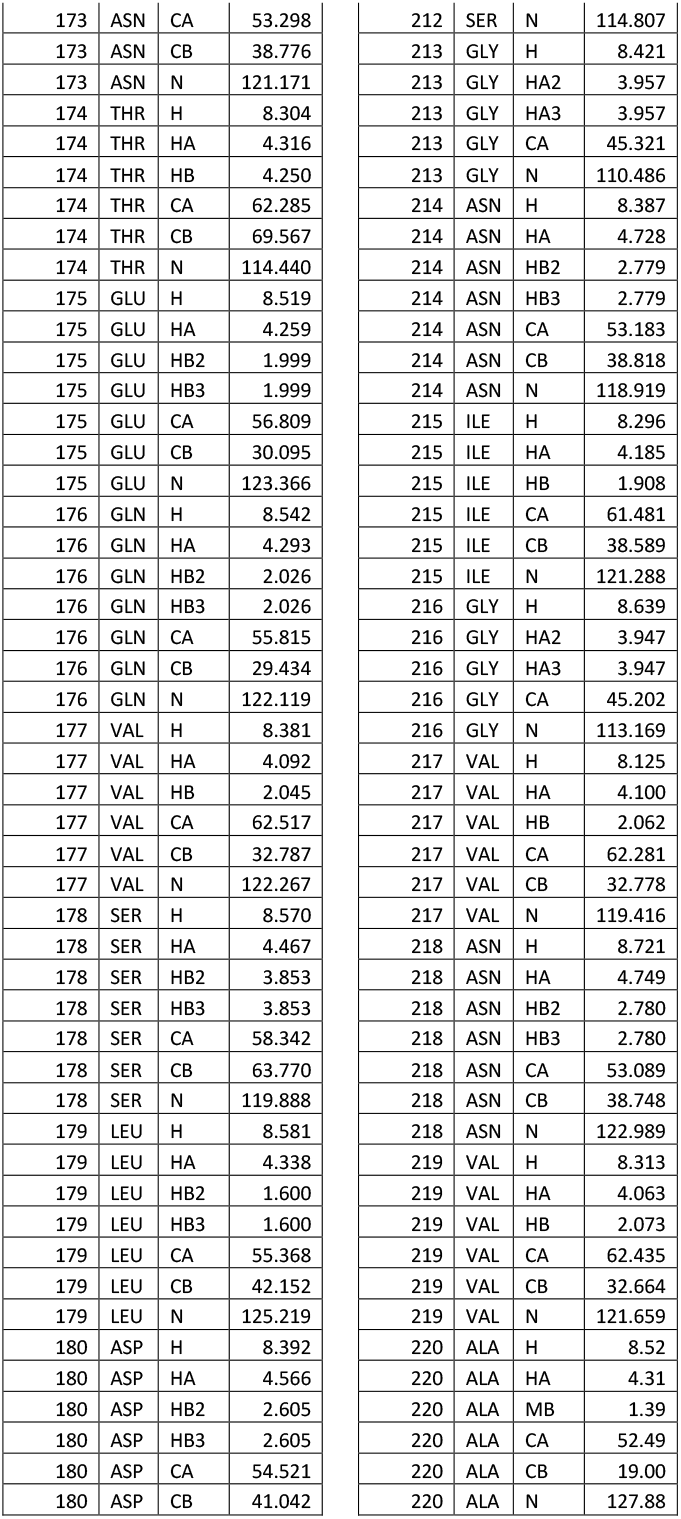
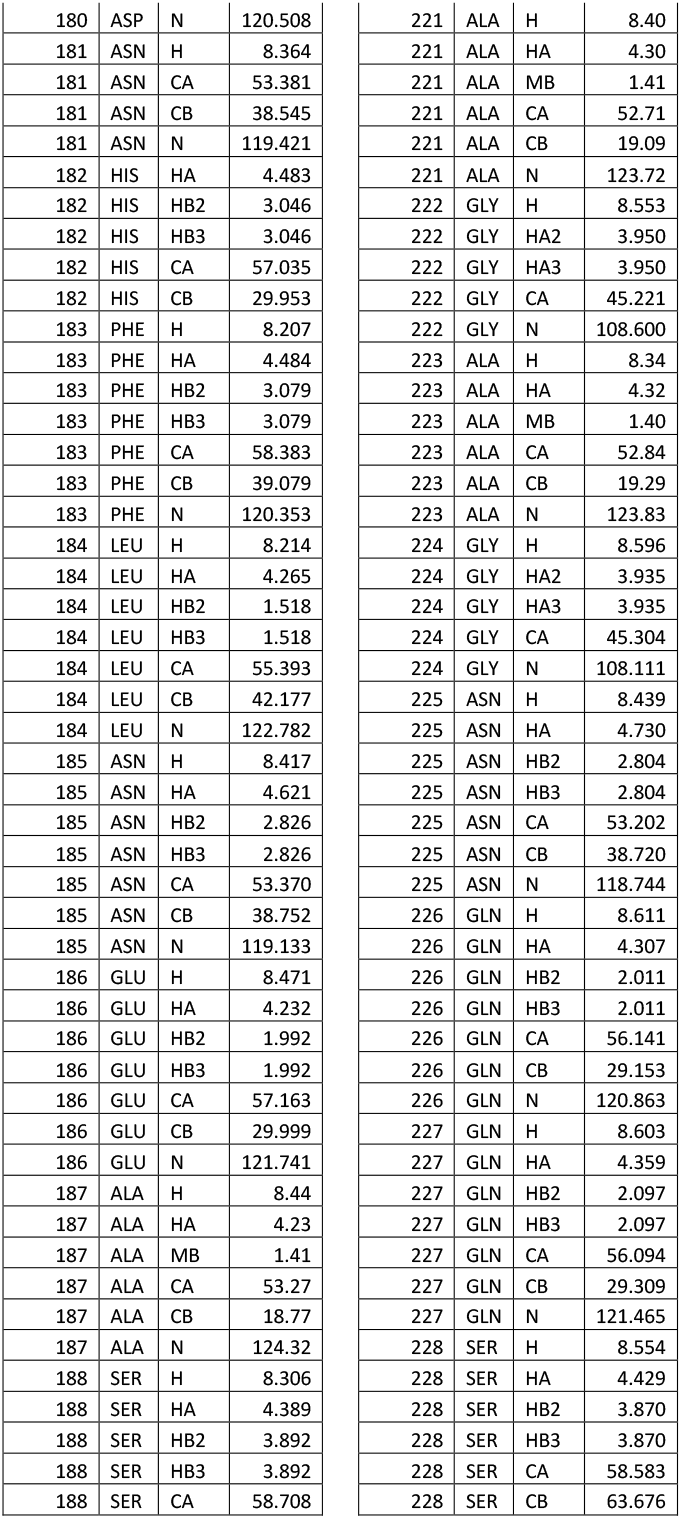
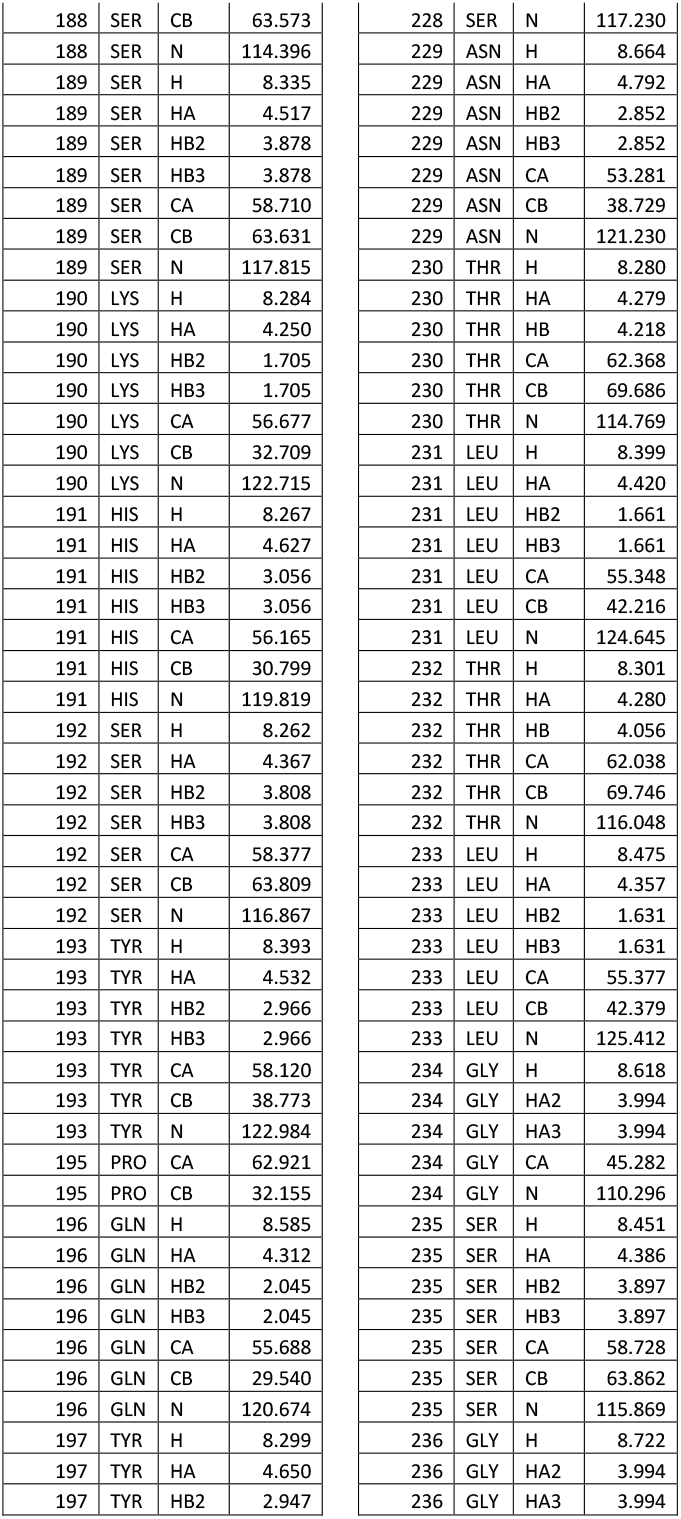
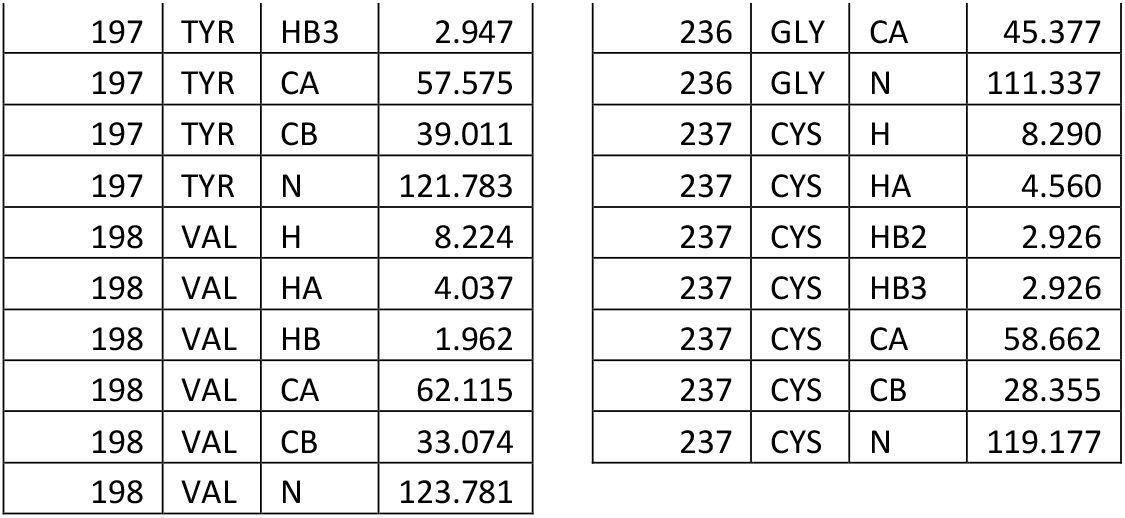
NMR resonance assignments of full-length FapC.

