## Supplementary figures and images for "Solution-state NMR Assignment and Secondary Structural Propensities of the Full-Length and Minimalistic-Truncated Prefibrillar Monomeric Form of Biofilm-Forming Functional-Amyloid FapC from *Pseudomonas aeruginosa*"

### Figure 1

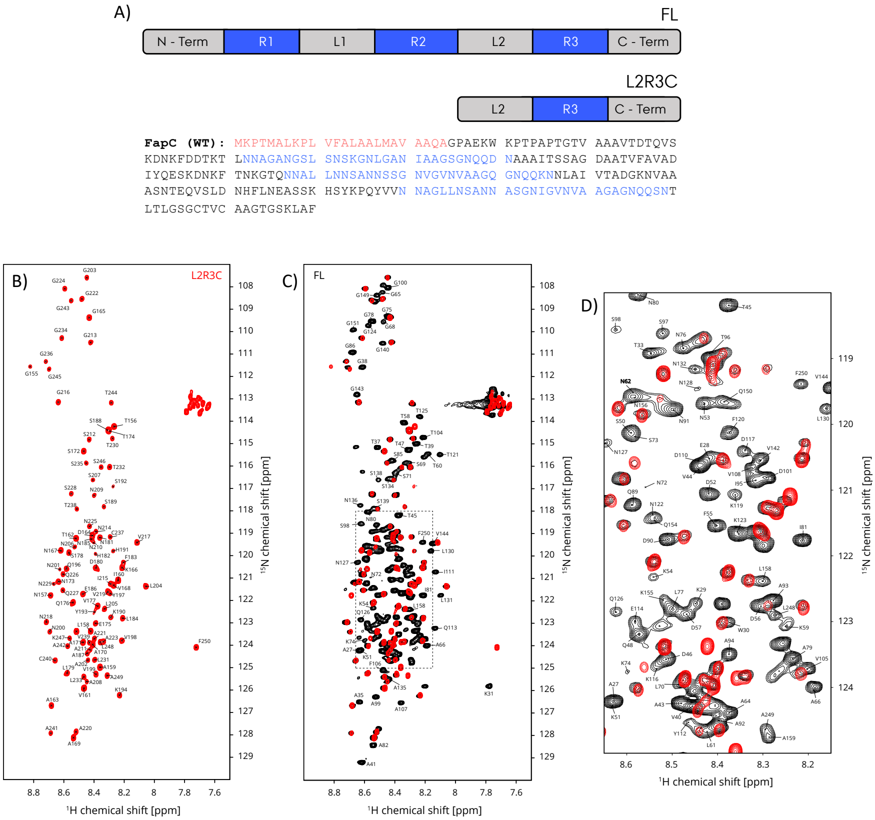

### Figure 2

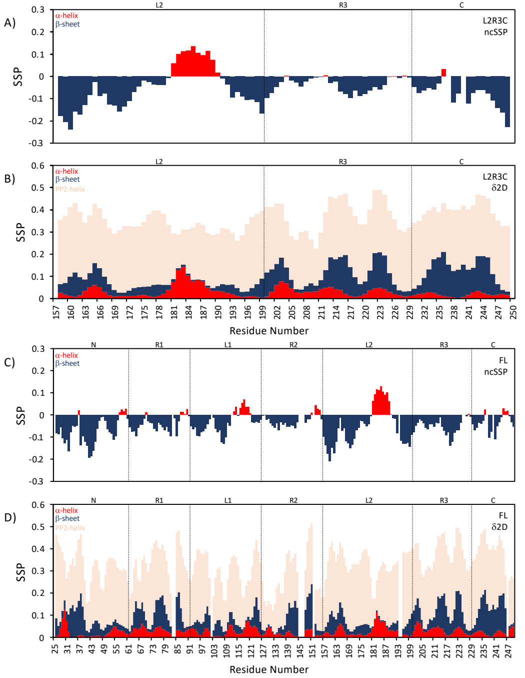

### Figure 3

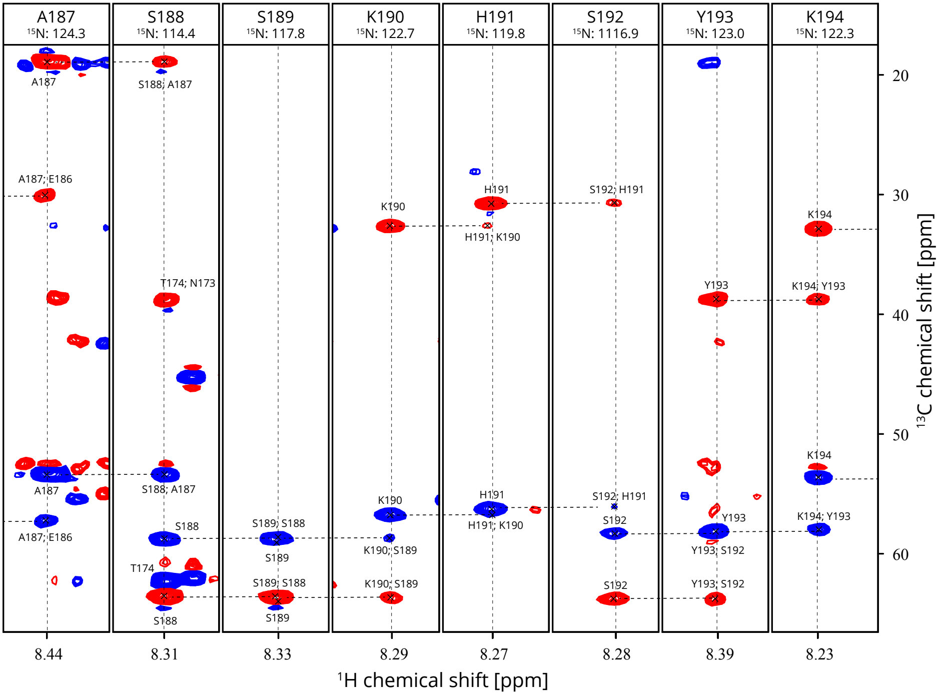
